# IF1 restrains excessive elevation of the mitochondrial membrane potential and safeguards the epithelial state of human iPSCs

**DOI:** 10.64898/2026.06.16.732339

**Authors:** Kaoru Kinjo, Gakuya Takamatsu, Kanako Toyama, Chitoshi Takayama, Yuko Akamine, Riko Kuniyoshi, Naoaki Otsuka, Yoko Manome, Hirotaka James Okano, Chiaki Katagiri, Mitsuyoshi Takatori, Masayuki Matsushita

## Abstract

Human pluripotent stem cells (hPSCs) rely on glycolysis and exhibit low mitochondrial respiration, conditions under which F1Fo ATP synthase tends to run in reverse, hydrolyzing ATP. ATP synthase inhibitory factor subunit 1 (IF1) inhibits F1Fo-mediated ATP hydrolysis, but its significance in glycolytic hPSCs remains unclear. Here, we show that stable IF1 knockdown (IF1-KD) in human induced pluripotent stem cells (hiPSCs) enhanced mitochondrial ATP hydrolysis and elevated the mitochondrial membrane potential (ΔΨm). Markers of the undifferentiated state were largely maintained, whereas IF1-KD cells exhibited reduced epithelial characteristics and a partial epithelial-mesenchymal transition (EMT)-like state. IF1-KD cells also showed enhanced store-operated Ca²⁺ entry (SOCE) and increased nuclear NFATc3. Pharmacological reduction of ΔΨm attenuated SOCE, whereas NFATc3 overexpression reproduced key features of the EMT-like expression pattern. Endogenous IF1 therefore acts as a constitutive restraint on reverse-mode F1Fo activity, limiting excessive ΔΨm and downstream SOCE–NFATc3 signaling and safeguarding the epithelial state of hPSCs.

**Graphical Abstract:** 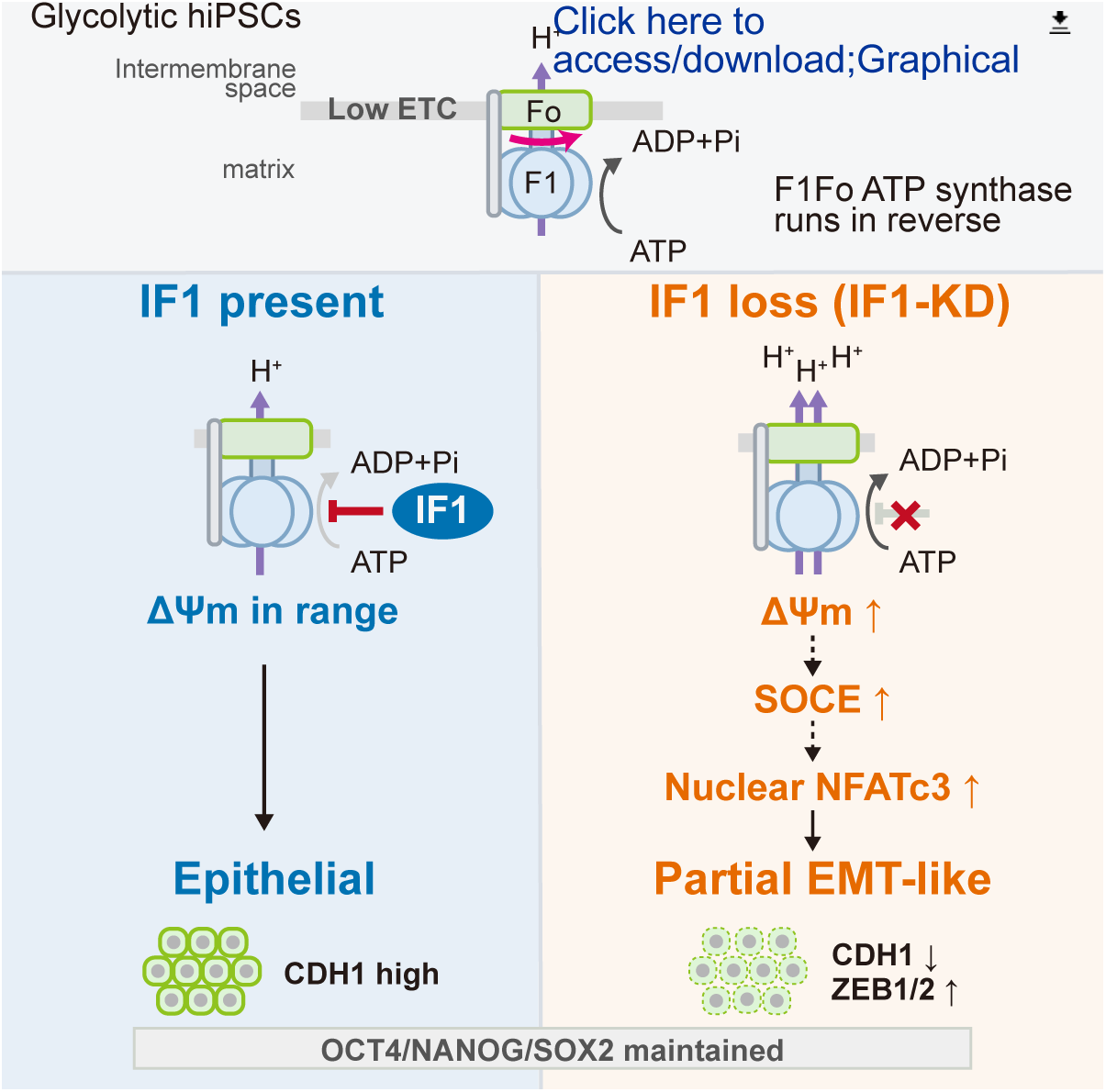

## Introduction

Human pluripotent stem cells (hPSCs), including embryonic stem cells (ESCs) and induced pluripotent stem cells (iPSCs), meet most of their energy demand through glycolysis and exhibit relatively low mitochondrial respiration (Varum et al., 2011; Zhang et al., 2011; Zhou et al., 2012). This glycolytic preference is not merely a passive adaptation to the hypoxic peri-implantation environment, but rather an intrinsic metabolic program that supports the pluripotent state(Teslaa and Teitell, 2015). By limiting mitochondria-derived reactive oxygen species (ROS), this metabolic configuration contributes to the maintenance of genomic stability (Saretzki et al., 2008). At the same time, glycolysis and partially maintained tricarboxylic acid (TCA) cycle flux provide precursors required to meet the high biosynthetic demand for nucleotides, amino acids, and lipids (Teslaa and Teitell, 2015), as well as metabolites that serve as substrates or cofactors for chromatin-modifying enzymes and thereby contribute to regulation of the PSC epigenetic state (Moussaieff et al., 2015; TeSlaa et al., 2016).

The electron transport chain (ETC) generates the proton motive force (pmf), comprising the mitochondrial membrane potential (ΔΨm) and a pH gradient, which drives ATP synthesis by F1Fo ATP synthase (hereafter F1Fo) (Junge and Nelson, 2015; Mitchell, 1961; 1966). F1Fo is reversible, however. When ETC function is compromised, as under hypoxic conditions, the pmf declines and F1Fo can operate in reverse, hydrolyzing ATP to pump protons and thereby support ΔΨm (Campanella et al., 2008). Although this reverse mode has traditionally been studied as an adaptive response to respiratory dysfunction (Campanella et al., 2009), recent work has shown that reverse F1Fo activity can also contribute to the pmf at steady state under glycolytic conditions (Rieger et al., 2021). hPSC mitochondria are typically globular with poorly developed cristae (St John et al., 2005; Varum *et al*., 2011), and their low ETC activity may favor constitutive operation of F1Fo in reverse. Consistent with this metabolic state, hPSCs produce ATP predominantly through glycolysis, and F1Fo has been proposed to contribute to ΔΨm maintenance by hydrolyzing glycolytically derived ATP (Zhang *et al*., 2011).

ATP synthase inhibitory factor subunit 1 (IF1), encoded by *ATP5IF1* (formerly *ATPIF1*), is a small mitochondrial matrix protein expressed in hPSCs (Vazquez-Martin et al., 2013). IF1 binds at an α/β catalytic interface of the F1 domain and selectively inhibits rotation of the γ subunit in the ATP-hydrolytic direction, thereby suppressing ATP hydrolysis by F1Fo (Cabezón et al., 2003; Gledhill et al., 2007; Pullman and Monroy, 1963). The net direction of the F1Fo reaction is determined by the pmf, the phosphorylation potential, and the extent of inhibition by IF1 (Junge and Nelson, 2015). IF1 has been studied primarily in the context of mitochondrial energy metabolism, first as a safeguard against futile ATP consumption when respiration is compromised, such as during ischemia or anoxia (Campanella *et al*., 2008), and later as a regulator of cancer-associated metabolic reprogramming (Formentini et al., 2012; Sánchez-Cenizo et al., 2010; Sgarbi et al., 2018b). More recent studies suggest that the physiological consequences of IF1 activity are highly dependent on cellular and tissue context (Brunetta et al., 2024; Domínguez-Zorita et al., 2023). However, how endogenous IF1 contributes to constitutive metabolism and maintenance of the cellular state in hPSCs remains unclear. If reverse-mode F1Fo activity is favored by the glycolytic, low-ETC state of hPSCs, IF1 may have an important role in maintaining the hPSC state by restraining this activity.

ΔΨm–Ca²⁺ signaling provides a potential route through which IF1-mediated control of F1Fo reverse mode could influence cell state. The undifferentiated hPSC state is characterized not only by the core pluripotency transcriptional network formed by OCT3/4, SOX2, and NANOG (Boyer et al., 2005), but also by a strong epithelial character (Li et al., 2012). This epithelial state is partially remodeled during differentiation and can involve epithelial–mesenchymal transition (EMT) (Thiery et al., 2009). EMT is not a binary switch, but encompasses a spectrum of intermediate hybrid states with both epithelial and mesenchymal features, often associated with increased cellular plasticity (Sha et al., 2019). Ca²⁺ signaling has been implicated in regulating such state transitions in PSCs. The calcineurin–nuclear factor of activated T-cells (NFAT) pathway promotes PSC differentiation, and NFATc3 has been reported to regulate lineage and EMT marker expression (Chen et al., 2021; Li et al., 2011). Store-operated Ca²⁺ entry (SOCE), a major Ca²⁺ influx pathway in non-excitable cells, provides sustained Ca²⁺ signals that promote calcineurin-dependent NFAT dephosphorylation and nuclear translocation (Hogan et al., 2003). ΔΨm, in turn, is the major driving force for mitochondrial Ca²⁺ uptake through the mitochondrial Ca²⁺ uniporter (MCU) (Baughman et al., 2011; De Stefani et al., 2011). By buffering Ca²⁺ entering through ORAI/calcium release-activated calcium (CRAC) channels, mitochondria can limit Ca²⁺-dependent channel inactivation and thereby sustain SOCE (Glitsch et al., 2002; Hoth et al., 2000). Whether IF1-mediated control of F1Fo reverse mode contributes to maintenance of the epithelial hPSC state through ΔΨm–Ca²⁺ signaling remains unknown.

Here, we investigated the role of IF1-mediated control of F1Fo reverse mode in hPSCs. We generated hiPSCs with stable IF1 knockdown (IF1-KD) and examined the consequences for mitochondrial function and cell state. IF1 depletion enhanced mitochondrial ATP hydrolytic activity and elevated ΔΨm, while largely preserving markers of the undifferentiated state. Nevertheless, IF1-KD cells exhibited reduced epithelial characteristics, accompanied by enhanced Ca²⁺–NFATc3 signaling. NFATc3 overexpression was sufficient to induce similar epithelial remodeling. These findings uncover a previously unrecognized connection between IF1-dependent mitochondrial regulation and maintenance of the epithelial state in hPSCs.

## Results

### F1Fo hydrolyzes ATP in reverse mode to maintain the mitochondrial membrane potential in hiPSCs

In bioenergetic states with low ETC activity, F1Fo tends to operate in reverse mode, hydrolyzing ATP to pump protons into the intermembrane space. hPSCs rely largely on glycolysis for energy production and have low ETC activity. Under these conditions, F1Fo has been reported to maintain the ΔΨm in part by hydrolyzing glycolytically derived ATP in hPSCs (Zhang *et al*., 2011).

To assess F1Fo ATP hydrolytic activity in hPSCs, we compared the hiPSC line 1383D6 with the human fetal lung fibroblast line TIG-1 and HeLa cells. ATP hydrolytic activity was measured in isolated mitochondria by an NADH-coupled spectrophotometric assay and normalized to total mitochondrial protein. hiPSCs showed significantly higher activity than both TIG-1 and HeLa cells (Figure 1A).

**Figure 1.**
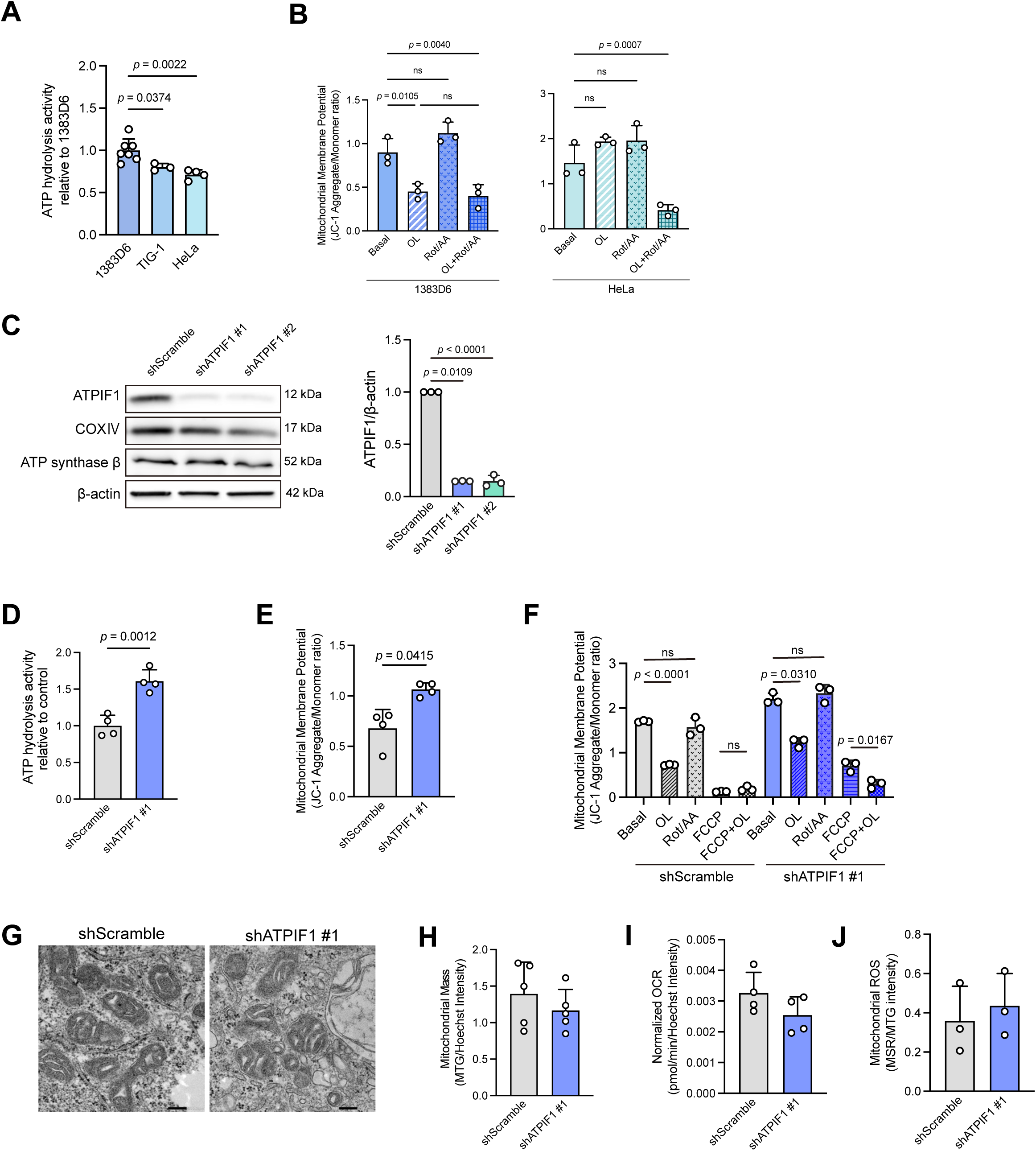
IF1 restrains F1Fo reverse mode and limits the mitochondrial membrane potential in hiPSCs. IF1 (ATP synthase inhibitory factor subunit 1) was knocked down with two independent lentiviral shRNAs, shATPIF1 #1 and #2. shScramble-transduced cells (SCR) served as the control, and shATPIF1 #1 is the representative line. IF1-overexpressing cells (ATPIF1-OE) were generated with a lentiviral vector and compared with empty vector. The mitochondrial membrane potential (ΔΨm) was assessed as the JC-1 aggregate/monomer fluorescence ratio in (B), (E) and (F). Panels (D–J) compare SCR and shATPIF1 #1 cells. (A) Representative quantification of ATP hydrolysis activity in isolated mitochondria from 1383D6 hiPSCs, TIG-1 fibroblasts, and HeLa cells, normalized to the mean of 1383D6 hiPSCs (set to 1). Seven independent experiments for 1383D6 hiPSCs and four each for TIG-1 and HeLa cells. (B) ÄØm in 1383D6 hiPSCs and HeLa cells treated with 10 μM oligomycin A (OL), an F1Fo inhibitor, or 2 μM each of rotenone and antimycin A (Rot/AA), respiratory chain inhibitors. Three independent experiments. (C) Representative western blot images of IF1, cytochrome c oxidase subunit Ⅳ (COX IV), ATP synthase F1 β subunit (ATP synthase β), and β-actin, and quantification of relative IF1 protein levels in shScramble, shATPIF1 #1 and shATPIF1 #2 cells, normalized to β-actin. Three independent experiments. IF1 protein is labeled as ATPIF1 in the figures. (D) Relative quantification of ATP hydrolysis activity in isolated mitochondria, normalized to the mean of the control group (set to 1). Four independent experiments. (E) ÄØm. Four independent experiments. (F) ÄØm in cells treated with 10 μM OL, 2 μM Rot/AA, 50 μM carbonyl cyanide 4-(trifluoromethoxy)phenylhydrazone (FCCP), or FCCP combined with OL (FCCP+OL). Three independent experiments. (G) Representative transmission electron microscopic images showing mitochondrial ultrastructure. Scale bar = 200 nm. (H) Quantification of mitochondrial mass. Five independent experiments. (I) Basal oxygen consumption rate (OCR), measured using an oxygen-sensitive probe-based assay. Four independent experiments. (J) Mitochondrial reactive oxygen species (mtROS) levels assessed by MitoSOX. Three independent experiments. Individual values with mean ± SD. Exact *P* values are shown when *P* < 0.05. Statistics: repeated-measures one-way ANOVA followed by Dunnett’s multiple comparison test (A, B), paired two-tailed t-tests on log-transformed values with Holm correction (C), unpaired two-tailed t-test (D), paired two-tailed t-test on log-transformed values (E, H–J), and repeated-measures two-way ANOVA followed by Bonferroni correction (F). See also Figure S1.

To determine whether this activity contributes to ΔΨm maintenance, we measured ΔΨm in cultured cells using the F1Fo inhibitor oligomycin A (OL). In hiPSCs, OL reduced the JC-1 ratio by approximately 50% relative to baseline (Figure 1B), indicating that F1Fo is required for ΔΨm maintenance. Localized F1Fo-mediated ATP hydrolysis has been reported in HeLa cells conditioned toward glycolysis in high-glucose medium (Rieger *et al*., 2021). In our hands, however, OL did not alter ΔΨm in HeLa cells (Figure 1B), indicating that the F1Fo dependence of ΔΨm is more pronounced in hiPSCs. To assess the contribution of the ETC in hiPSCs, we added the complex I/III inhibitor rotenone/antimycin A (Rot/AA), which did not significantly alter ΔΨm (Figure 1B). Together, these results indicate that the ΔΨm in hiPSCs depends largely on F1Fo rather than on the ETC.

### Loss of IF1 enhances F1Fo reverse mode and elevates the mitochondrial membrane potential in hiPSCs

To examine whether IF1 regulates F1Fo-mediated ATP hydrolysis in hiPSCs, we generated stable IF1-KD hiPSCs using a lentiviral shRNA system. To exclude off-target effects, two independent lines were established with distinct shRNA sequences (shATPIF1 #1 and shATPIF1 #2), with shScramble-transduced (SCR) cells as controls. Western blotting confirmed that IF1 protein levels were reduced by more than 80% (Figure 1C). The expression of cytochrome c oxidase subunit IV (COX IV) and the ATP synthase F1 β subunit (ATP synthase β) was unchanged, indicating that neither mitochondrial mass nor F1Fo abundance was affected. In parallel, we generated IF1-overexpressing (IF1-OE) cells, in which IF1 protein was increased 5-fold with no change in COX IV or ATP synthase β (Figure S1B).

We then measured ATP hydrolytic activity in isolated mitochondria; activity was significantly increased approximately 1.6-fold in IF1-KD cells relative to SCR (Figure 1D). The JC-1 ratio was significantly elevated 1.6-fold in IF1-KD cells (Figure 1E). A comparable elevation was observed in shATPIF1 #2, although the limited number of independent experiments (n = 2) precluded statistical evaluation (Figure S1A).

Conversely, IF1 overexpression significantly reduced ATP hydrolytic activity to 85% of control (Figure S1C) without altering the basal ΔΨm (Figure S1D). OL nonetheless reduced ΔΨm in IF1-OE cells (Figure S1D), indicating that F1Fo reverse mode remains active and that increased IF1 abundance alone does not abolish it.

To determine the source of ΔΨm in IF1-KD cells, we measured ΔΨm in cells pretreated with OL or Rot/AA. In both SCR and IF1-KD cells, OL markedly reduced ΔΨm, whereas Rot/AA had no significant effect (Figure 1F). This pattern mirrored that of parental hiPSCs (Figure 1B), indicating that ΔΨm in these cells is sustained primarily by F1Fo reverse mode rather than by respiration. However, the extent of OL-induced reduction was comparable between the two lines and thus did not by itself demonstrate enhanced reverse mode. ΔΨm in the presence of OL remained higher in IF1-KD than in SCR cells, a difference that our assay could not resolve further.

We therefore assessed reverse mode under uncoupling with carbonyl cyanide 4-(trifluoromethoxy)phenylhydrazone (FCCP). Because reverse-mode flux is constrained by the prevailing ΔΨm itself, uncoupling relieves this constraint and reveals differences in reverse-mode capacity. In SCR cells, FCCP alone was sufficient to dissipate ΔΨm almost completely, and co-treatment with FCCP and OL produced no further change (Figure 1F). In IF1-KD cells, by contrast, FCCP alone left a substantial residual ΔΨm, and near-complete dissipation required the combination of FCCP and OL (Figure 1F). Thus, in IF1-KD cells F1Fo continues to hydrolyze ATP and pump protons even when the proton gradient is dissipated.

Together with the increased ATP hydrolytic capacity of isolated mitochondria (Figure 1D), these results indicate that endogenous IF1 constitutively restrains F1Fo reverse mode in hiPSCs.

### IF1 depletion elevates ΔΨm without activating respiration or perturbing net energy metabolism

Although Rot/AA did not reduce ΔΨm in either SCR or IF1-KD cells (Figure 1F), suggesting that the ETC does not contribute to ΔΨm formation, we further examined whether mitochondrial respiration is activated in IF1-KD cells, both morphologically and functionally. Transmission electron microscopic analysis showed sparse, underdeveloped cristae in both SCR and IF1-KD cells (Figure 1G). Mitochondrial mass, assessed by MitoTracker Green staining, did not differ between the two groups (Figure 1H). Basal oxygen consumption rate (OCR) was comparable between SCR and both IF1-KD lines (Figures 1I and S1E). Mitochondrial ROS (mtROS) did not differ between groups (Figure 1J). These results indicate that the elevated ΔΨm in IF1-KD cells reflects enhanced F1Fo ATP hydrolysis rather than increased mitochondrial respiration.

Finally, we examined the effect of enhanced F1Fo ATP hydrolysis on cellular energy metabolism. Basal intracellular ATP levels did not differ between SCR and IF1-KD cells (Figure S1F). After OL addition, intracellular ATP increased by approximately 25% in both groups, consistent with the cessation of ATP hydrolysis, although this change was not statistically significant (Figure S1F). Glucose consumption and lactate production were also unchanged in both KD lines (Figure S1G). These results indicate that the ATP flux through F1Fo reverse mode is small relative to whole-cell ATP turnover, and that the enhanced ATP hydrolysis caused by IF1 knockdown has only a minor effect on intracellular ATP levels and on the balance between glycolysis and oxidative phosphorylation.

### IF1 depletion impairs epithelial character while preserving the undifferentiated state in hiPSCs

Because IF1 depletion elevated the ΔΨm without perturbing energy metabolism, we next examined whether IF1-depleted cells differ in their properties as hiPSCs.

The mRNA levels of the undifferentiated-state markers *POU5F1* (OCT4), *NANOG*, and *SOX2* did not differ between SCR and IF1-KD cells, and immunocytochemistry for SSEA4 and NANOG confirmed that the undifferentiated state was maintained (Figures 2A and 2B). Proliferation rate was also not significantly altered (Figure S2A). These results indicate that IF1-KD cells retain self-renewal and the undifferentiated state.

**Figure 2.**
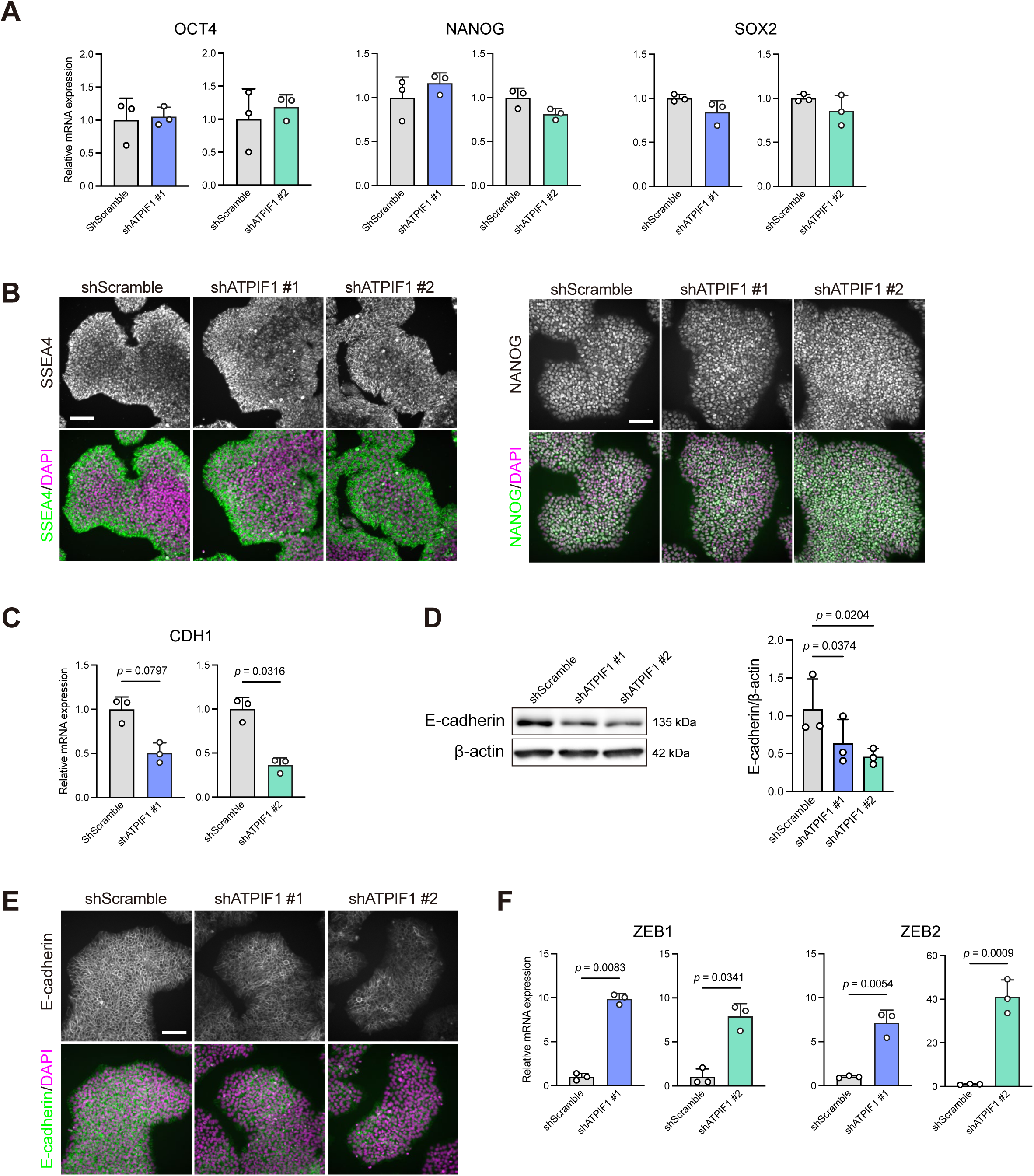
IF1 depletion reduces E-cadherin and induces *ZEB1* and *ZEB2* while preserving markers of the undifferentiated state in hiPSCs. All panels compare shScramble, shATPIF1 #1, and shATPIF1 #2 cells. RT-qPCR data were normalized to *GAPDH* and expressed relative to the mean of the corresponding control group (set to 1). Three independent experiments each. (A) RT-qPCR of the undifferentiated-state markers *POU5F1* (OCT4), *NANOG*, and *SOX2*. (B) Representative immunocytochemistry images of stage-specific embryonic antigen-4 (SSEA4, green) and NANOG (green), each with DAPI (magenta). Scale bar = 100 μm. (C) RT-qPCR of *CDH1*. (D) Representative western blot images of E-cadherin and quantification of relative E-cadherin protein levels, normalized to β-actin. (E) Representative immunocytochemistry images of E-cadherin (green) and DAPI (magenta). Scale bar = 100 μm. (F) RT-qPCR of the epithelial-mesenchymal transition (EMT)-related transcription factors *ZEB1* and *ZEB2*. Individual values with mean ± SD. Exact *P* values are shown when *P* < 0.05. Statistics: paired two-tailed t-test on log-transformed values (A, C, F) and repeated-measures one-way ANOVA followed by Dunnett’s multiple comparison test (D). See also Figure S2.

We next examined epithelial character, another hallmark of undifferentiated hPSCs. *CDH1* mRNA, which is abundantly expressed in PSCs (Li *et al*., 2012), showed a decreasing tendency that varied between the two KD lines, whereas E-cadherin protein was consistently and significantly decreased in both lines (Figures 2C and 2D). Immunocytochemistry further showed reduced membrane-localized E-cadherin signal in IF1-KD cells, confirming the decrease in E-cadherin at both the protein and localization levels (Figure 2E). Because loss of E-cadherin is a hallmark of EMT, we examined the expression of EMT-related genes by RT-qPCR. The EMT-inducing transcription factors *ZEB1* and *ZEB2* were significantly increased in both IF1-KD lines (Figure 2F). To confirm that this change is reproducible across hiPSC lines, we generated IF1-KD cells in an independent line, jri1401#15, which showed the same expression changes, namely reduced *CDH1* and increased *ZEB1* (Figure S2B). In contrast, IF1-OE did not significantly alter the expression of these genes (Figure S2C). Together, these results indicate that IF1 depletion preserves self-renewal and the expression of OCT4, NANOG, and SOX2, while selectively impairing the epithelial character associated with the undifferentiated state.

### IF1 depletion induces a partial EMT-like state and activates calcium–NFATc3 signaling

To clarify what transcriptional changes underlie the selective loss of epithelial character in IF1-KD cells, we performed RNA-seq on SCR and shATPIF1 #1 cells. Principal component analysis (PCA) separated the two groups along PC1 (Figure 3A). The EMT-inducing transcription factors *ZEB1* and *ZEB2* were the top contributors to PC1, followed by developmental genes (*PITX2, LMO3, TFAP2A, UNC5C*) and pluripotency-associated genes (*ZFP42, HHLA1*) (Figure 3B). The core pluripotency factors *POU5F1*, *NANOG*, and *SOX2*, however, were not among the top contributors, consistent with their unchanged expression (Figure 2A).

**Figure 3.**
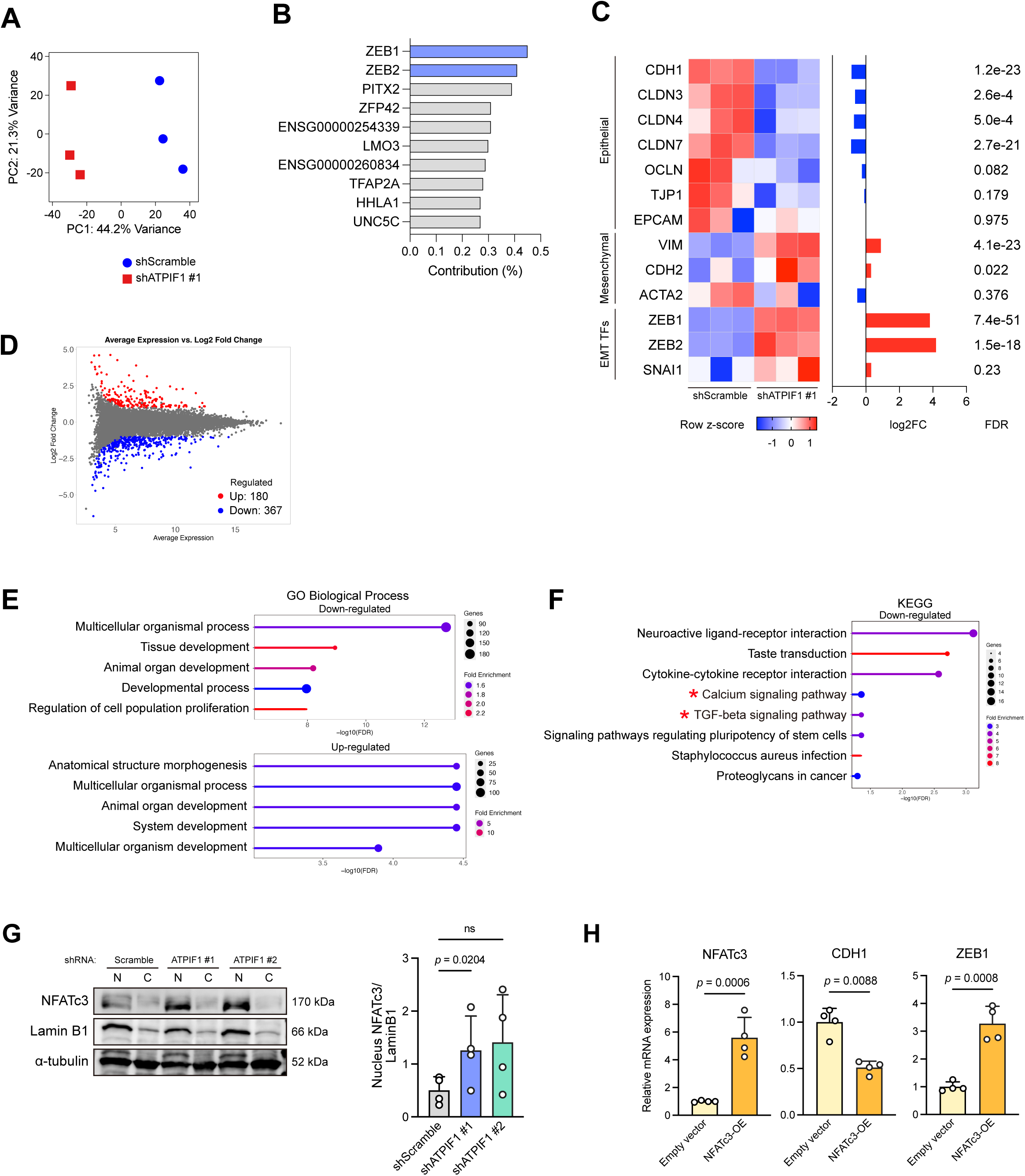
IF1-KD hiPSCs show a partial EMT-like signature accompanied by NFATc3 nuclear translocation. (A–F) RNA sequencing of shScramble and shATPIF1 #1 cells, with three independent experiments per group. Differentially expressed gene (DEG) identification with DESeq2, principal component analysis (PCA), and enrichment analyses were performed using iDEP version 2.4.4. (A) PCA of shScramble and shATPIF1 #1 cells. PC1 and PC2 account for 44.2% and 21.3% of the total variance, respectively. (B) Top 10 genes contributing to PC1. (C) Expression of epithelial markers, mesenchymal markers, and EMT-related transcription factors (EMT TFs) in shScramble and shATPIF1 #1 cells. The heatmap shows row z-scores of log2(TPM + 1); log2 fold change and FDR from DESeq2 are shown at right. Genes with baseMean < 30 (*SNAI2*, *TWIST1*, *TWIST2*) were excluded owing to low expression. Three independent experiments. (D) MA plot of DEGs (FDR < 0.1 and |log2FC| > 1). Upregulated and downregulated genes are shown in red and blue, respectively. (E, F) Gene Ontology (GO) biological process (E) and Kyoto Encyclopedia of Genes and Genomes (KEGG) pathway (F) enrichment analyses. In (E), the top five enriched GO terms are shown for each of the upregulated and downregulated DEGs. In (F), the top eight enriched KEGG pathways among the downregulated DEGs are shown. The x-axis indicates −log10(FDR), dot size indicates the number of genes, and dot color indicates fold enrichment. (G) Western blots and quantification of nuclear factor of activated T cells, cytoplasmic 3 (NFATc3), normalized to Lamin B1, in shScramble, shATPIF1 #1, and shATPIF1 #2 cells. Four independent experiments. (H) RT-qPCR of *NFATc3*, *CDH1*, and *ZEB1* in empty vector and NFATc3-overexpression (NFATc3-OE) cells, normalized to *ATP5PB* and expressed relative to the mean of the control group (set to 1). Four independent experiments. Data in (G) and (H): individual values with mean ± SD. Exact *P* values are shown when *P* < 0.05. Statistics: repeated-measures one-way ANOVA followed by Dunnett’s multiple comparison test (G) and paired two-tailed t-test on log-transformed values (H). See also Figure S3.

We next examined a curated set of epithelial markers, mesenchymal markers, and EMT transcription factors using the DESeq2 results (Figure 3C, Table S1). Epithelial markers were selectively reduced in IF1-KD cells: *CDH1* and the claudins *CLDN3*, *CLDN4*, and *CLDN7* were significantly downregulated, whereas *EPCAM* and *TJP1* were unchanged despite comparable expression levels. Among mesenchymal markers, *VIM* and *CDH2* were upregulated whereas ACTA2 was unchanged. Among EMT transcription factors, *ZEB1* and *ZEB2* were strongly induced, whereas *SNAI1* was unchanged. *SNAI2, TWIST1, and TWIST2* were excluded owing to low expression (Figure 3C). IF1-KD cells therefore appear to occupy a partial EMT-like state, in which selective activation of the *ZEB1* and *ZEB2* and loss of junctional epithelial markers occur without induction of *SNAI1* or of mesenchymal structural genes.

To assess transcriptional changes genome-wide, we identified differentially expressed genes (DEGs) using DESeq2 (FDR < 0.1, |log2 fold change| > 1), detecting 180 upregulated and 367 downregulated genes (Figure 3D). Gene Ontology (GO) biological process analysis revealed terms related to development, proliferation, and differentiation among both upregulated and downregulated DEGs, suggesting that IF1-KD broadly affects developmental and differentiation processes (Figure 3E). Moreover, Kyoto Encyclopedia of Genes and Genomes (KEGG) analysis revealed enrichment of signaling pathways among the downregulated DEGs, including the calcium and TGF-β signaling pathways (FDR = 0.0493 for both; Figure 3F).

We next examined canonical EMT-inducing pathways: phosphorylation of SMAD2/3 (TGF-β/Activin), ERK1/2 (FGF), and GSK-3β (Wnt), and nuclear translocation of β-catenin (Wnt). None differed significantly between SCR and IF1-KD lines (Figure S3), suggesting that these pathways are unlikely to account for the partial EMT-like state.

Among the pathways identified by KEGG analysis, calcium signaling remained a candidate. However, transcriptional enrichment alone does not indicate the direction of signaling, as the pathway includes both positive and negative regulators. To assess the direction directly, we examined NFAT transcription factors, which are activated by calcium signaling and have been implicated in cell plasticity and differentiation (Hogan *et al*., 2003). NFATc3 has been reported to induce EMT in PSCs (Chen *et al*., 2021; Li *et al*., 2011). Nuclear translocation of NFATc3 was significantly increased in shATPIF1 #1 cells, and showed a similar trend in shATPIF1 #2 (Figure 3G). We therefore overexpressed NFATc3 in hiPSCs (jri line); NFATc3-overexpressing (NFATc3-OE) cells showed decreased *CDH1* and increased *ZEB1* (Figure 3H), matching the pattern in IF1-KD cells (Figure 2C, F). These results suggest that calcium–NFATc3 signaling is activated in IF1-KD cells and that NFATc3 activation is sufficient to induce key features of the partial EMT-like state.

### Elevated mitochondrial membrane potential enhances store-operated Ca²⁺ entry in IF1-depleted hiPSCs

To understand how calcium–NFATc3 signaling is activated in IF1-KD cells, we examined whether the elevated ΔΨm (Figure 1) shapes intracellular Ca²⁺ dynamics. We performed ratiometric Ca²⁺ imaging and found that basal intracellular Ca²⁺ levels did not differ between groups (Figure 4A). To assess SOCE, we depleted endoplasmic reticulum (ER) Ca²⁺ stores with thapsigargin (TPG) and monitored Ca²⁺ influx. IF1-KD cells showed a significantly higher Ca²⁺ peak than SCR cells, with the peak ΔR/R₀ approximately twice that of SCR cells (Figures 4B and 4C). The rise was also faster in IF1-KD cells, with a shorter time to peak (Figure 4D). In the late phase, however, the Ca²⁺ level at 300 s was significantly lower in IF1-KD than in SCR cells (Figure 4E).

**Figure 4.**
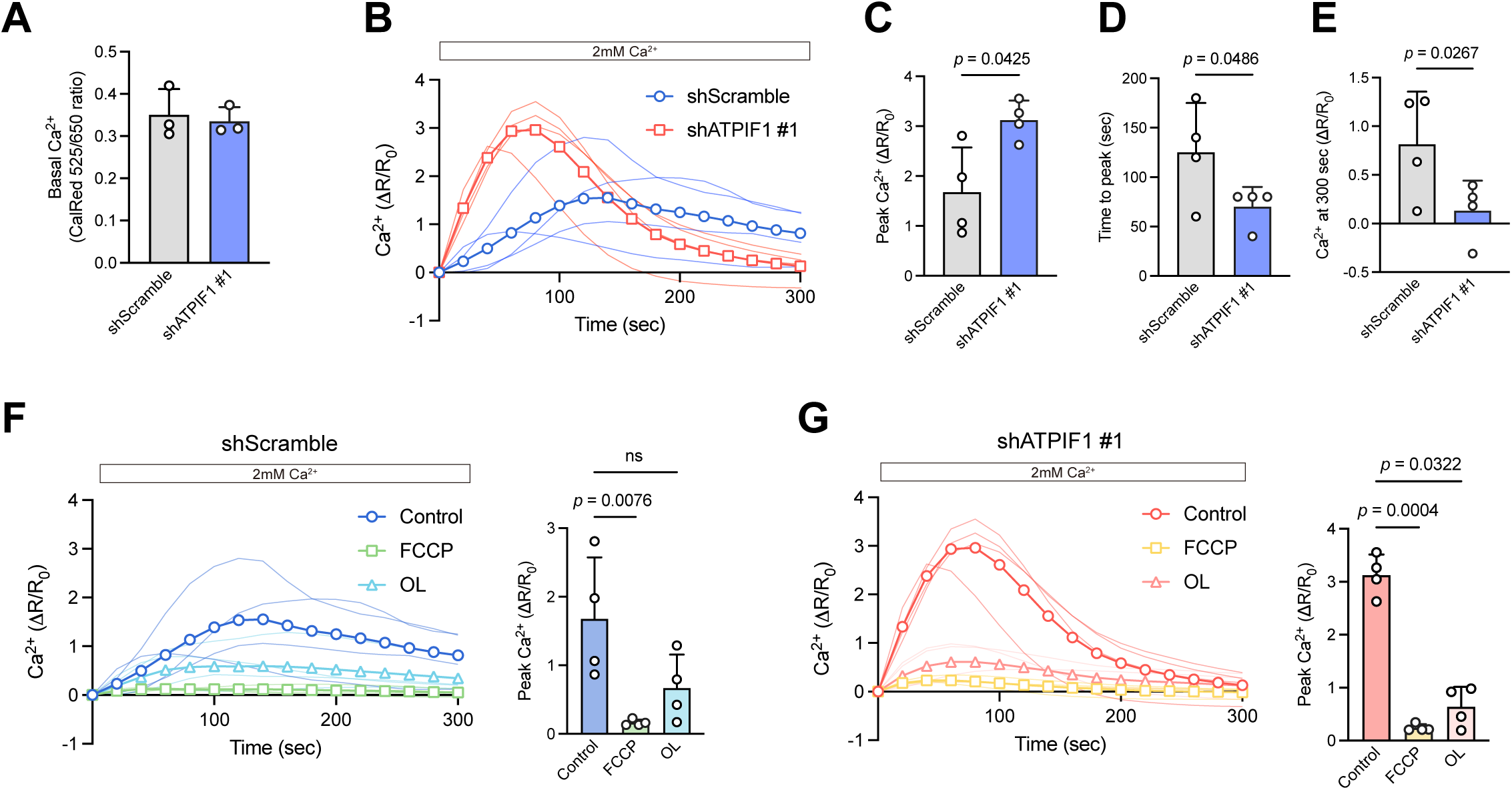
IF1 depletion enhances ΔΨm-dependent store-operated Ca²⁺ entry in hiPSCs. All panels compare shScramble and shATPIF1 #1 cells. Four independent experiments, except (A), which comprises three. (A) Basal intracellular Ca²⁺ levels (CalRed 525/650 ratio). (B) Traces of store-operated Ca²⁺ entry (SOCE), recorded after re-addition of 2 mM CaCl₂ at t = 0 s to thapsigargin (TPG)-treated cells in Ca²⁺-free buffer. Bold lines indicate the mean and thin lines individual experiments. (C–E) Peak SOCE amplitude (C), time to peak (D), and Ca²⁺ level at 300 s (E), derived from (B). (F, G) SOCE traces and peak amplitudes in shScramble (F) and shATPIF1 #1 (G) cells under control conditions (no inhibitor) and in the presence of 50 μM FCCP or 10 μM OL. Traces were recorded as in (B). The control traces are those in (B), replotted for comparison. Individual values are shown with the mean ± SD. Exact *P* values are shown when *P* < 0.05. Statistics: paired two-tailed t-test on log-transformed values (A, C–E) and repeated-measures one-way ANOVA followed by Dunnett’s multiple comparison test (F, G). See also Figure S4.

We next tested the ΔΨm dependence of this response. Dissipating the ΔΨm with the uncoupler FCCP almost completely abolished SOCE in both SCR and KD cells (Figures 4F and 4G). This indicates that mitochondrial Ca²⁺ uptake driven by the ΔΨm is essential for the maintenance of SOCE in hiPSCs, as previously reported in other cell types (Glitsch *et al*., 2002; Hoth *et al*., 2000). Furthermore, the addition of OL significantly reduced the SOCE peak to approximately 50–70% of the respective controls in both SCR and IF1-KD cells (Figures 4F and 4G), consistent with the OL-induced reduction in ΔΨm to about half of the basal level shown in Figure 1. These results indicate that SOCE in hiPSCs depends on the ΔΨm and that lowering the ΔΨm attenuates SOCE.

To determine whether the enhanced SOCE could be explained by transcriptional changes, we examined the expression of ten genes regulating SOCE and mitochondrial Ca²⁺ handling in the RNA-seq data, including *ORAI1*, *STIM1*, *MCU*, and voltage-dependent anion channel 1 (*VDAC1*) (Figure S4A). Only *VDAC1*, an outer-membrane channel mediating mitochondrial Ca²⁺ uptake, was significantly downregulated in IF1-KD cells. Because reduced *VDAC1* would act to suppress rather than enhance SOCE, the elevated SOCE in IF1-KD cells cannot be explained by *VDAC1* expression. *MCU* transcript levels were unchanged. However, MCU protein has been reported to increase upon IF1 depletion in HeLa cells (Faccenda et al., 2021). We therefore examined MCU by western blotting and found no difference between SCR and IF1-KD cells (Figure S4B). Together, these results suggest that the elevated ΔΨm, rather than altered expression of SOCE regulators, is the principal factor underlying the enhanced SOCE in IF1-KD cells.

### The EMT-like state persists once established, independently of continued upstream signaling

The preceding results indicate that IF1-KD elevates the ΔΨm, enhances SOCE, and activates NFATc3, and that NFATc3 activation is sufficient to induce EMT-like gene expression. We next asked whether the EMT-like state, once established, remains dependent on this axis.

We targeted the axis at two levels: downstream, by inhibiting NFAT signaling with the calcineurin inhibitor cyclosporin A (CsA), and upstream, by blocking the ΔΨm-driven mitochondrial Ca²⁺ uptake that sustains SOCE with the MCU inhibitor Ru265. Under the conditions examined, neither CsA nor Ru265 restored CDH1 or ZEB1 expression in IF1-KD cells (Figures 5A and 5B). These findings do not exclude a role for the ΔΨm–SOCE–NFAT axis in establishing the EMT-like state, but indicate that the established state is not readily reversed by subsequent inhibition of the axis.

**Figure 5.**
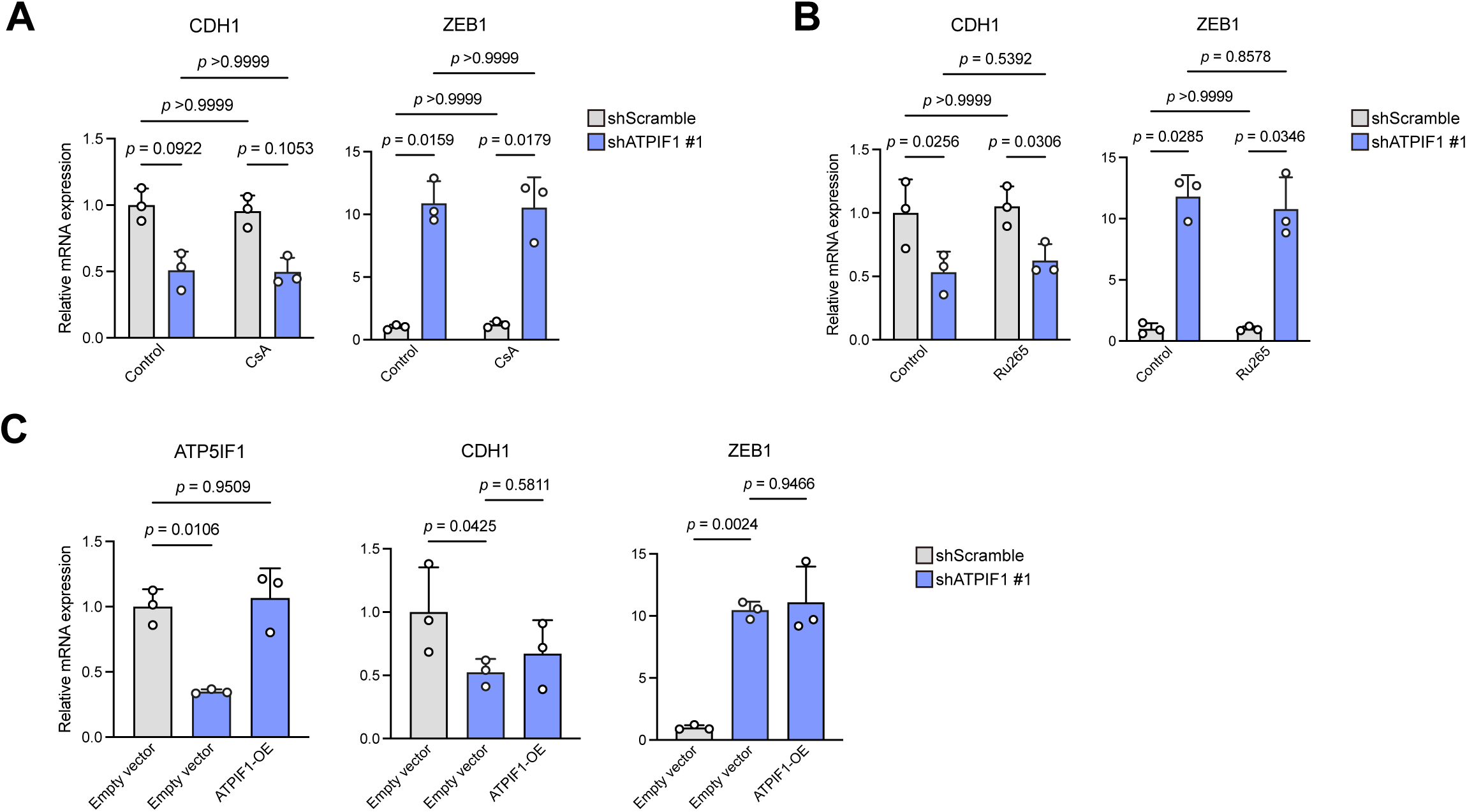
The established EMT-like state in IF1-KD hiPSCs is not reversed by calcineurin or MCU inhibition or by *ATP5IF1* re-expression. All panels compare shScramble and shATPIF1 #1 cells. RT-qPCR data were normalized to *ACTB* and expressed relative to the mean of the corresponding control group (set to 1). Three independent experiments. (A) RT-qPCR of *CDH1* and *ZEB1* after treatment for 24 h with 1 μM cyclosporin A (CsA), an inhibitor of calcineurin. DMSO served as the vehicle control. (B) RT-qPCR of *CDH1* and *ZEB1* after treatment for 24 h with 30 μM Ru265, an inhibitor of the mitochondrial Ca²⁺ uniporter (MCU). Untreated cells served as the control. (C) RT-qPCR of *ATP5IF1*, *CDH1*, and *ZEB1* in IF1-KD cells transduced with a lentiviral IF1-OE vector. Individual values with mean ± SD. Exact *P* values are shown for the comparisons of interest, including those that did not reach statistical significance. Statistics: repeated-measures two-way ANOVA followed by Bonferroni correction (A, B), repeated-measures one-way ANOVA followed by Dunnett’s multiple comparison test (C).

Finally, we re-expressed IF1 in IF1-KD cells to determine whether restoration of IF1 could reverse the EMT-like state. Although *ATP5IF1* mRNA expression was restored, no clear recovery of CDH1 or ZEB1 expression was observed (Figure 5C). Together, these results indicate that, once established, the EMT-like state of IF1-KD cells is not readily reversed by late inhibition of calcineurin or MCU, or by restoration of IF1 expression.

### IF1 depletion promotes mesodermal differentiation with EMT while suppressing endodermal and neuroectodermal differentiation

A pre-existing partial EMT-like state may alter responses to differentiation cues. We therefore examined trilineage differentiation.

To induce mesendodermal differentiation, we activated Wnt signaling by inhibiting GSK-3 with CHIR99021 (Figure 6A). At 24 h after induction, more migrating cells were observed at the colony edges in IF1-KD than in SCR cells (Figures 6B and S5A). We assessed the cadherin switch, a hallmark of EMT, by immunocytochemistry. Even before induction, the N-cadherin/E-cadherin ratio was higher in IF1-KD than in SCR cells (Figure S5B), consistent with Figure 2. At 24 h, this ratio was unchanged in SCR but significantly increased in IF1-KD cells (Figure 6C), indicating that the cadherin switch, already evident before induction, was further promoted upon differentiation. In a time-course RT-qPCR analysis, the mesendoderm/primitive streak marker *TBXT* (brachyury) showed a trend toward upregulation in IF1-KD cells that did not reach significance owing to large variability, whereas the early mesoderm marker *TBX6* was significantly upregulated at 33 h (Figure 6D). The mesendoderm subsequently bifurcates into mesoderm and endoderm. When we induced definitive endoderm differentiation with Activin A (Figure 6A), the mRNA levels of the endodermal markers *FOXA2* and *SOX17* were significantly lower in IF1-KD cells after induction (Figure 6E).

**Figure 6.**
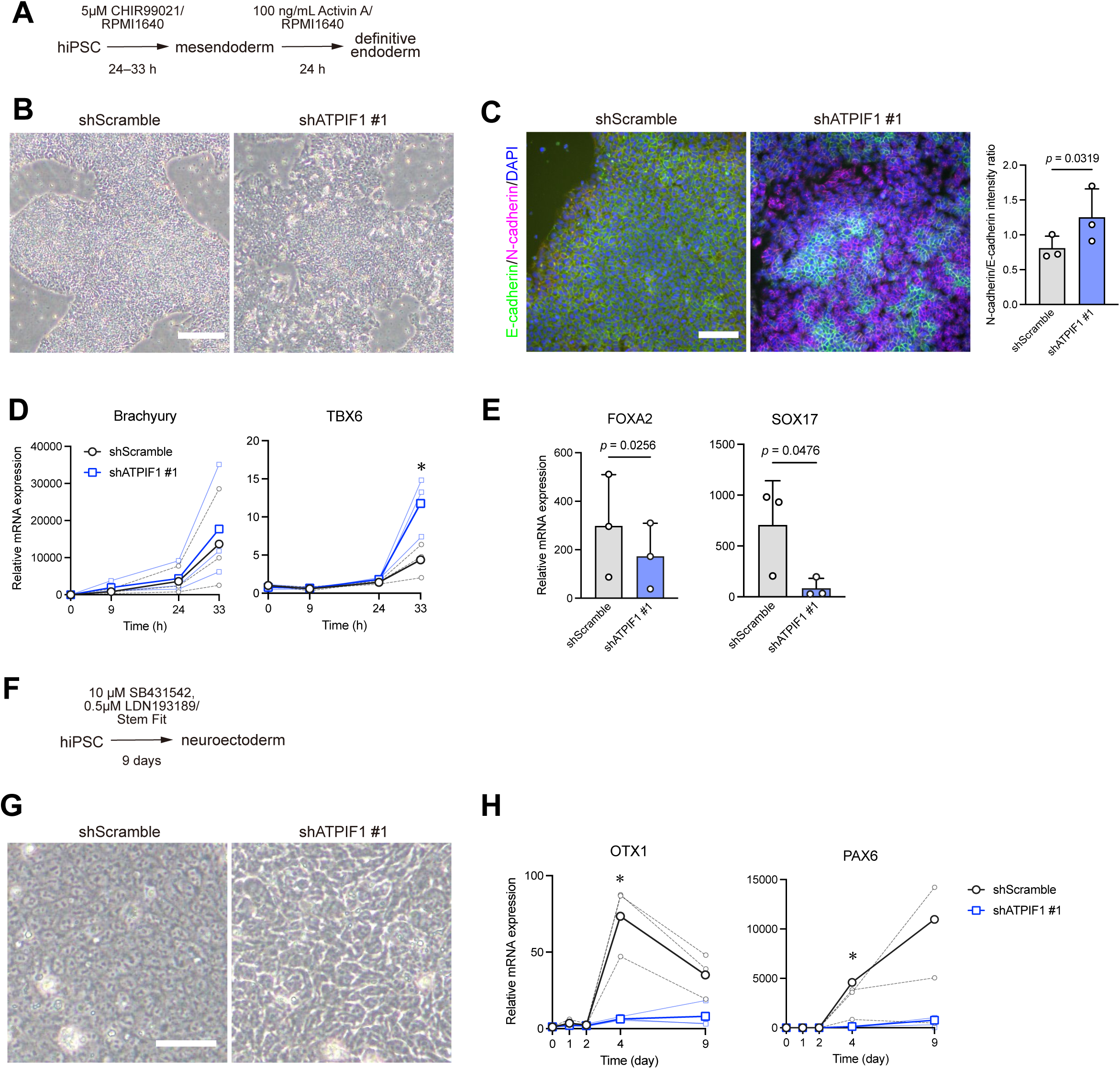
IF1 depletion promotes mesodermal differentiation while suppressing endodermal and neuroectodermal differentiation in hiPSCs. All panels compare shScramble and shATPIF1 #1 cells. RT-qPCR data were normalized to *ACTB* and expressed as a fold-change relative to shScramble cells at 0 h (set to 1). Three independent experiments. (A) Schematic of the differentiation protocol for mesendoderm and definitive endoderm. (B) Representative phase-contrast images 24 h after mesendodermal differentiation induction. Scale bar = 200 μm. (C) Representative immunocytochemistry images of E-cadherin (green), N-cadherin (magenta), and DAPI (blue), and quantification of the N-cadherin/E-cadherin intensity ratio at 24 h after mesendodermal differentiation induction. Scale bar = 100 μm. (D) Time-course RT-qPCR of the mesendoderm/mesoderm markers *TBXT* (Brachyury) and *TBX6*. Bold lines indicate the mean, while thin lines indicate individual experiments. (E) RT-qPCR of the definitive endoderm markers *FOXA2* and *SOX17* at day 2 of endoderm induction. (F) Schematic of the dual-SMAD inhibition (2Si) protocol for neuroectoderm induction. (G) Representative phase-contrast images 2 days after 2Si. Scale bar = 50 μm. (H) Time-course RT-qPCR of the neuroectoderm markers *OTX1* and *PAX6*. Bold lines indicate the mean, while thin lines indicate individual experiments. Data in (C–E) and (H): individual values with mean ± SD. *Exact *P* values are shown where indicated; *P* < 0.05. Statistics: paired two-tailed t-test on log-transformed values, applied at 33 h in (D) and at day 4 in (H). See also Figure S5.

We next performed dual SMAD inhibition to induce neuroectodermal differentiation (Figure 6F). By day 2 of induction, SCR cells had already formed a dense, epithelial-like sheet, whereas IF1-KD cells remained enlarged and failed to form a comparable dense structure through day 9, the last time point examined (Figures 6G and S5C). Consistently, the neuroectodermal marker *OTX1* rose approximately 73-fold at day 4 relative to day 0 in SCR but only 6-fold in IF1-KD cells (Figure 6H). *PAX6* expression was also significantly lower in IF1-KD at day 4 and remained low through day 9 (Figure 6H). Notably, the mesendodermal/mesodermal, endodermal, and neuroectodermal markers did not differ between SCR and IF1-KD cells before differentiation (Figure S5D). Together, these findings indicate that IF1 depletion alters differentiation responses, favoring mesoderm-associated changes while attenuating endodermal and neuroectodermal programs.

## Discussion

Our study identifies endogenous IF1 as a constitutive restraint on F1Fo reverse-mode activity in glycolytic hiPSCs, where its loss reduces epithelial identity without compromising the core pluripotency network. Loss of IF1 increased the ATP hydrolytic activity of mitochondria and raised ΔΨm. Although IF1-KD cells largely maintained undifferentiated-state markers, they showed reduced E-cadherin expression, a partial EMT-like transcriptional state, enhanced SOCE, and increased nuclear accumulation of NFATc3. Two experiments address causality at distinct steps of this chain: pharmacological reduction of ΔΨm attenuated SOCE, and NFATc3 overexpression alone reproduced key features of the EMT-like expression pattern. Together, these findings support a model in which IF1 tunes reverse-mode F1Fo activity to set the resting ΔΨm, thereby limiting downstream SOCE–NFATc3 signaling and preventing the onset of epithelial remodeling in undifferentiated hiPSCs.

Our findings support the operation of F1Fo in reverse mode in hiPSCs under basal conditions. Compared with somatic TIG-1 cells and cancer-derived HeLa cells, hiPSCs exhibited significantly higher mitochondrial ATP hydrolytic activity, and OL lowered ΔΨm. These findings indicate that reverse-mode F1Fo makes a substantial contribution to ΔΨm maintenance in hiPSCs, supporting the proposal by Zhang et al. that glycolytically derived ATP contributes to ΔΨm maintenance in these cells (Zhang *et al*., 2011).

Our results further indicate that the physiologically consequential output of IF1 is the control of ΔΨm rather than the conservation of cellular ATP. Because reverse-mode F1Fo couples ATP hydrolysis to proton pumping, IF1 affects both processes. The consequences, however, differ greatly in magnitude. IF1 depletion did not measurably alter intracellular ATP levels, glucose consumption, or lactate production, suggesting that ATP flux through reverse-mode F1Fo is small relative to whole-cell ATP turnover. ΔΨm, by contrast, was markedly elevated, and this elevation remained sensitive to OL but not to Rot/AA. Thus, IF1 modulates ΔΨm without substantially altering net cellular energy metabolism in undifferentiated hiPSCs.

The IF1-overexpression experiments provide further insight into how IF1 regulates reverse-mode activity. IF1 overexpression reduced ATP hydrolytic activity in isolated mitochondria but did not alter basal ΔΨm in intact cells. One possible explanation is that the inhibitory activity of IF1 depends not only on its abundance but also on its regulatory state, including pH-dependent oligomerization and protein kinase A-mediated phosphorylation (Cabezon et al., 2000; García-Bermúdez et al., 2015). Thus, increasing IF1 abundance may not necessarily increase its effective inhibitory activity under basal conditions. Moreover, OL continued to reduce ΔΨm in IF1-OE cells, indicating that reverse-mode F1Fo remained active despite increased IF1 expression. Together, these observations are consistent with a model in which IF1 tunes rather than abolishes reverse-mode activity. Whether IF1 also inhibits ATP synthesis under conditions supporting oxidative phosphorylation remains debated (Carroll et al., 2024; García-Bermúdez *et al*., 2015; Sánchez-Cenizo *et al*., 2010). This question is unlikely to be central in hiPSCs, in which the ETC contributes little to ΔΨm maintenance, but it may become relevant as mitochondrial metabolism is remodeled during differentiation.

Our findings suggest that the elevated ΔΨm contributes to the altered SOCE observed in IF1-KD cells. SOCE in these cells was characterized by a higher and more rapidly rising peak, followed by a faster decline and lower late-phase cytosolic Ca²⁺ levels. This response was strongly ΔΨm-sensitive: FCCP nearly abolished SOCE, whereas OL attenuated the SOCE peak to comparable levels in SCR and IF1-KD cells, supporting a functional link between mitochondrial polarization and SOCE in hiPSCs. One possible mechanism is enhanced MCU-mediated mitochondrial Ca²⁺ uptake driven by the elevated ΔΨm. Mitochondrial Ca²⁺ buffering can limit Ca²⁺-dependent inactivation of CRAC channels and sustain Ca²⁺ influx, while simultaneously accelerating cytosolic Ca²⁺ clearance (Hoth *et al*., 2000; Hoth et al., 1997; Naghdi et al., 2010). Such a mechanism could account for both the larger SOCE peak and the more rapid decay observed in IF1-KD cells. The reduced late-phase Ca²⁺ signal might appear inconsistent with the classical view that NFAT activation requires a sustained low-amplitude Ca²⁺ plateau (Dolmetsch et al., 1997). However, NFAT can be preferentially activated by local Ca²⁺ microdomains near open CRAC channels rather than by the spatially averaged cytosolic signal (Kar et al., 2011), so the larger and faster SOCE response may still generate a stronger local signal for NFAT activation. Notably, IF1 depletion has been reported to increase MCU expression through AMPK/CREB signaling in HeLa cells (Faccenda *et al*., 2021), whereas MCU protein abundance was unchanged in IF1-KD hiPSCs. Thus, enhanced SOCE cannot be explained by increased MCU abundance and is instead consistent with altered mitochondrial Ca²⁺ handling associated with elevated ΔΨm.

Our findings further implicate Ca²⁺–NFATc3 signaling in the partial EMT-like state. Nuclear accumulation of NFATc3 increased in IF1-KD cells, and NFATc3 overexpression reproduced key features of the IF1-KD transcriptional phenotype, including reduced *CDH1* and increased *ZEB1* expression, consistent with previous reports that NFAT signaling promotes EMT-associated programs in PSCs (Chen *et al*., 2021; Li *et al*., 2011). Interestingly, partial EMT induced by *CDH1* knockdown in hPSCs has been reported to revert when the knockdown is released (Aban et al., 2021). In contrast, the EMT-like state established in IF1-KD cells was not reversed by inhibition of calcineurin with cyclosporin A or MCU with Ru265, nor by restoration of IF1 expression. These findings suggest that NFATc3 contributes to establishing the EMT-like state, which, once established, becomes persistent and less dependent on continued upstream signaling while undifferentiated-state markers remain largely maintained.

Our data also indicate that the mitochondrial signal linked to the EMT-like state is more closely associated with ΔΨm than with mtROS. IF1 depletion altered neither cristae morphology, mitochondrial mass, basal oxygen consumption, nor mtROS levels, in contrast to somatic and cancer cells in which IF1 has been linked to changes in mitochondrial respiration, cristae organization, and mtROS production (Faccenda *et al*., 2021; Formentini *et al*., 2012; Romero-Carramiñana et al., 2023). A high ΔΨm can favor reverse electron transport (RET) and thereby increase mtROS production. However, mtROS did not increase in IF1-KD cells. Importantly, the higher ΔΨm in these cells would tend to increase mitochondrial MitoSOX accumulation rather than obscure an increase in mtROS, making it unlikely that enhanced mtROS accounts for the EMT-like phenotype. The lack of mtROS elevation may reflect the low respiratory activity of hiPSCs, which provides limited electron flux to support RET. Consistent with this, no ROS-related terms were significantly enriched in the GO or KEGG analyses. Together, these findings suggest that in glycolytic hiPSCs, altered mitochondrial polarization itself, rather than increased mtROS, may contribute to changes in epithelial state.

This altered epithelial state was also accompanied by changes in the response to differentiation cues. IF1-KD cells showed a stronger mesoderm-associated response during mesendoderm induction, but weaker responses to definitive endoderm and neuroectoderm differentiation. The enhanced mesoderm-associated response may be related to the pre-existing partial EMT-like state, because mesoderm formation involves loss of epithelial organization and acquisition of migratory properties (Thiery *et al*., 2009). By contrast, our data do not distinguish whether the reduced endodermal and neuroectodermal responses reflect altered lineage choice or secondary consequences of the disrupted epithelial state. Mitochondria are increasingly recognized as regulators of developmental transitions (Granath-Panelo and Kajimura, 2024; Lemma et al., 2026; Schieke et al., 2008), but whether IF1 participates in such physiological regulation remains unresolved. Because mitochondrial metabolism is rapidly and lineage-specifically remodeled during differentiation (Folmes et al., 2012), the bioenergetic context supporting the mechanism described here is likely to shift accordingly. Addressing this will require temporally controlled IF1 depletion together with analysis of IF1 expression in developing organisms.

Our study has several limitations. Although pharmacological reduction of ΔΨm attenuated SOCE, we did not test the converse, namely whether increasing ΔΨm is sufficient to enhance SOCE. Addressing this will require an approach that elevates ΔΨm independently of ETC activity and ATP synthesis. We also did not determine whether lowering ΔΨm can restore epithelial marker expression once the EMT-like state has been established. Although inhibition of calcineurin or MCU and re-expression of IF1 also failed to reverse the state, the efficacy of these interventions was not directly verified. In addition, mitochondrial Ca²⁺ uptake, which we propose contributes to the altered SOCE response, was not measured directly. Although we found no evidence for activation of canonical EMT-associated pathways, including TGF-β, Wnt, and FGF signaling, or for increased mtROS, we cannot exclude contributions from non-mitochondrial ROS sources such as NADPH oxidases or from other signaling pathways. A recent study showing that elevated ΔΨm can promote nuclear DNA methylation through mitochondrial membrane phospholipid remodeling (Mori et al., 2025) further raises the possibility that additional ΔΨm-dependent mechanisms operate in parallel with the Ca²⁺–NFAT pathway described here.

The evidence for reverse-mode F1Fo activity is itself indirect, resting on ATP hydrolysis in isolated mitochondria together with consistent pharmacological responses. Under FCCP-induced uncoupling, near-complete dissipation of ΔΨm in IF1-KD cells required the additional presence of OL, consistent with continued F1Fo-mediated proton pumping despite uncoupling. A similar persistence has been reported in IF1-deficient cancer cells (Sgarbi et al., 2018a). An unresolved observation is that ΔΨm remained higher in IF1-KD than in SCR cells even after OL treatment. Because Rot/AA had no detectable effect, respiration is unlikely to account for this residual ΔΨm; one possibility is incomplete inhibition of the elevated F1Fo activity by OL, although we cannot exclude a lower inner-membrane proton leak in IF1-KD cells, including an ANT-mediated component (Brand et al., 2005). A further consideration is the JC-1 signal responds non-linearly to ΔΨm; the magnitude of the differences reported here, including this residual signal, should therefore be interpreted as relative rather than proportional to ΔΨm in millivolts. Measurements with a potentiometric probe that responds linearly to ΔΨm, together with mitochondria-targeted ATP and pH sensors, will be needed to resolve reverse-mode F1Fo activity in living hiPSCs.

Notwithstanding these limitations, our findings point to a requirement imposed by the metabolic configuration of hPSCs. The glycolytic, low-ETC state of hPSCs has generally been considered advantageous for pluripotency because it limits mitochondria-derived ROS and supplies precursors and metabolites for biosynthesis and epigenetic regulation (Saretzki *et al*., 2008; Teslaa and Teitell, 2015). This same configuration, however, favors reverse-mode operation of F1Fo, so that ΔΨm is sustained by hydrolysis of glycolytically derived ATP. Reverse-mode F1Fo activity therefore creates a need to restrain excessive ΔΨm — a requirement absent in cells that maintain ΔΨm through respiration. In IF1-KD cells, ΔΨm elevation was accompanied by a partial EMT-like state, whereas markers of the undifferentiated state were largely preserved. In undifferentiated hiPSCs, endogenous IF1 may act as a safeguard against excessive ΔΨm, allowing reverse-mode F1Fo to contribute to ΔΨm maintenance while protecting the epithelial state.

## Resource availability

### Lead contact

Further information and requests for resources and reagents should be directed to and will be fulfilled by the lead contact, Gakuya Takamatsu.

### Materials availability

This study did not generate new unique reagents.

### Data and code availability

The data reported in the study are available from the lead contact upon reasonable request. The RNA-seq datasets generated in this study will be deposited in Japanese Genotype-phenotype Archive (JGA) for controlled-access raw sequence data (accession numbers will be assigned) prior to publication. This study did not generate new code.

## Acknowledgements

We would like to thank all of our collaborators and laboratory members for their support. We thank Dr. D. Utsumi for building the environment for bcbio-nextgen. We are grateful to I. Ikemiya for secretarial assistance. We thank Dr. H. Uchima for performing the earlier experiments that led to this study. We are grateful for helpful discussions with Dr. F. Wei, Dr. S. Li, Dr. C. Toma, and Dr. M. Nakamura. 1383D6 (HPS1006) was provided by the RIKEN BRC through the National BioResource Project of the MEXT, Japan. Cells were established by Kyoto University from Cellular Technology Limited (http://www.immunospot.com/) ePBMC® (Nakagawa et al., 2014). pLKO.1-blast and pLKO.1-blast-SCRAMBLE were gifts from Keith Mostov (Addgene plasmid # 26655 and # 26701).

This study was funded by the Sasakawa Scientific Research Grant from The Japan Science Society; JSPS Grant-in-Aid for Scientific Research [Grant Numbers 23K07017, 26K09850, 26K10494]; the advanced medical research grant in the Faculty of Medicine, University of the Ryukyus; the 6th Life Science Project from the Seimei Igaku Kenkyu Shinko Foundation. The funders had no role in the design and conduct of the study and the decision to submit the manuscript for publication.

## Author contributions

K.K., G.T., and M.M. conceived and designed the study. K.K., G.T., K.T., Y.A., M.T., and C.T. developed the methodology. K.K. and G.T. performed the formal analysis. K.K., G.T., K.T., Y.A., R.K., N.O., and C.T. carried out the investigation. K.K., K.T., and Y.A. curated the data. Y.M. and H.J.O. provided resources. K.K. wrote the original draft, and G.T., N.O., C.K., M.T., C.T., and M.M. reviewed and edited the manuscript. K.K. prepared the visualization. G.T., C.K., M.T., C.T., and M.M. supervised the study. G.T. and M.M. administered the project. K.K., G.T., M.T., and M.M. acquired funding. All authors agree to be accountable for all aspects of the work in ensuring that questions related to the accuracy or integrity of any part of the work are appropriately investigated and resolved. The corresponding author has full access to all the data in the study and is responsible for the decision to submit for publication.

## Declaration of interests

The authors declare that they have no known competing financial interests or personal relationships that could have appeared to influence the work reported in this paper.

## EXPERIMENTAL MODEL AND STUDY PARTICIPANT DETAILS

### Human induced pluripotent stem cell lines

The hiPSC line 1383D6 (RRID: CVCL_UP39, hPSCreg: BRCi014-A) was provided by the RIKEN BRC through the National BioResource Project of the Ministry of Education, Culture, Sports, Science and Technology (MEXT), Japan. The line had been established at Kyoto University using ePBMC® from Cellular Technology Limited (http://www.immunospot.com/) derived from a healthy 36-year-old male donor of Asian ancestry (Nakagawa et al., 2014), using episomal vectors. The cells were obtained at passage 14. The hiPSC line jri1401#15 (unique ID: not registered) was established from peripheral blood mononuclear cells of a healthy Japanese male in his 60s, using episomal vectors by the Jikei University School of Medicine, as previously described (Tajiri et al., 2018). The cells were obtained at passage 36.

Both lines were maintained under feeder-free conditions in StemFit AK02N medium (Ajinomoto) on iMatrix-511 (Takara) according to the manufacturer’s instructions. Cells were dissociated with TrypLE Select CTS (Thermo Fisher Scientific) diluted to 0.5× in 0.5 mM EDTA/PBS and passaged every 3–4 days at a density of 2 × 10² cells/mm² in the presence of 10 μM Y-27632 (FUJIFILM Wako). Cells were cryopreserved in STEM-CELLBANKER (Takara) and stored in liquid nitrogen. Thawed cells were plated in StemFit AK02N supplemented with 10 μM Y-27632. All cells were cultured at 37°C in 5% CO₂. Parental hiPSCs were used within 10 passages after receipt. The identities of both parental lines were based on the documentation provided by the supplying institutions; the lines were not independently re-authenticated in this study. Normal karyotypes of the parental hiPSC lines have been reported previously. All hiPSC lines used in this study tested negative for mycoplasma contamination using a PCR-based detection kit (Takara). All cultures were maintained without antibiotics, and cultures were monitored for bacterial and fungal contamination at every passage. Formal bacteriostasis and fungistasis testing was not performed.

### Genetically modified hiPSC lines

IF1-knockdown (shATPIF1 #1 and shATPIF1 #2), shScramble control, IF1-overexpressing (IF1-OE), and empty vector control lines were generated by lentiviral transduction followed by blasticidin selection, as described in the method details. These cells were maintained as bulk (non-clonal) populations without single-cell cloning, cryopreserved, and subsequently used for experiments. All stable transduced lines were derived from 1383D6. In addition, shScramble and shATPIF1 #1 lines were independently generated from jri1401#15 (Figure S2B).

For each stable transduced line, the passage at which cells were cryopreserved following completion of blasticidin selection was designated as passage 0 for subsequent experiments. The corresponding original passage numbers were P21 for shScramble and shATPIF1 #1, P24 for shATPIF1 #2, P22 for empty vector and IF1-OE, and P44 for shScramble and shATPIF1 #1 derived from jri1401#15. shScramble and shATPIF1 #1 cells derived from 1383D6 were used for up to 15 passages after cryopreservation, whereas all other stably transduced lines were used for up to 10 passages.

Karyotyping was performed on shScramble, shATPIF1 #1, and shATPIF1 #2 cells at their original passage numbers of P30, P30, and P32, respectively, by Nihon Gene Research Laboratories, Inc. using G-banding analysis of 20 metaphase cells. All three lines showed a normal 46, XY karyotype.

The undifferentiated state of the shScramble, shATPIF1 #1, and shATPIF1 #2 lines was confirmed by expression of POU5F1, NANOG, and SOX2 and by immunostaining for NANOG and SSEA4 (Figures 2A and 2B). Trilineage differentiation capacity was assessed by directed differentiation toward ectodermal, mesodermal, and endodermal lineages, followed by RT-qPCR analysis of two lineage-specific markers for each germ layer (Figure 6). All assessments were performed using cells within the passage range used for the experiments.

### Other cell lines

The human fetal lung-derived fibroblast-like cell line TIG-1 (RRID: CVCL_3181), derived from a female fetus at 20 weeks of gestation(Ohashi et al., 1980), was provided by the JCRB Cell Bank and was cultured in MEM (Thermo Fisher Scientific) supplemented with 10% fetal bovine serum and 1% GlutaMAX; cells were used at population doubling levels between 32 and 40. HeLa cells (female) were from an in-house laboratory stock of unknown passage history and were cultured in DMEM supplemented with 10% fetal bovine serum. LentiX-293T cells (Takara; female), used for lentiviral packaging, were cultured in DMEM supplemented with 10% fetal bovine serum. All cells were maintained at 37°C in 5% CO₂, and all tested negative for mycoplasma contamination using a PCR-based detection kit.

### Ethics statement

This study was approved by the Research Ethics Committee of the University of the Ryukyus (approval number 714) and the Research Ethics Committee of the Jikei University School of Medicine (approval number 27-083(7968)), and was conducted in accordance with the Declaration of Helsinki. hiPSCs were established from a healthy donor, who provided written informed consent for the establishment and research use of the cells.

This study used only established hiPSC lines cultured in vitro and did not involve human embryos, gametes or embryo models. It therefore corresponds to category 1A research under the ISSCR Guidelines for Stem Cell Research and Clinical Translation (2021) and was conducted in accordance with the ISSCR Standards for Human Stem Cell Use in Research (2023).

## METHOD DETAILS

### Lentiviral transduction for gene knockdown and overexpression

To generate hiPSC lines with IF1-KD, IF1-OE, or NFATc3-OE, lentiviral transduction was performed. For knockdown, duplex oligonucleotides encoding shRNA targeting human *ATP5IF1* (RefSeq: NM_016311.5) were cloned into the pLKO.1-blast vector (Addgene, #26655). The target sequence for shATPIF1 #1 and shATPIF1 #2 were derived from The RNAi Consortium (Broad Institute, TRCN0000149949) and a previous report (Fujikawa et al., 2012), respectively. For overexpression, a custom lentiviral vector, pLVSIN-EF1α-blast, was constructed by replacing the CMV promoter and a puromycin resistance gene in pLVSIN-CMV-Puro (Takara, #6183) with the EF1α promoter and a blasticidin resistance gene, respectively. Full-length coding sequences of human *ATP5IF1* or *NFATc3* (RefSeq: NM_173163.3) were cloned into this vector using the In-Fusion HD Cloning Kit (Takara, #638947). pLKO.1-blast-SCRAMBLE (Addgene, #26701) or pLVSIN-EF1α-blast (empty) were used as control vectors. Lentiviral particles were produced in LentiX-293T cells (Takara, #632180) with Lentiviral High Titer Packaging Mix (Takara, #6194) and Trans IT-Lenti Transfection Reagent (Mirus Bio, #MIR6600) following the manufacturer’s protocols. hiPSCs were transduced at a multiplicity of infection (MOI) of 10 (IF1-OE and NFATc3-OE) or 30 (IF1-KD). For establishment of stable cell lines of IF1-KD or IF1-OE, cells were cultured with 5 µg/mL blasticidin for 12 days and cryopreserved. Stable IF1-KD lines were generated from 1383D6 hiPSCs using two independent shRNAs (shATPIF1 #1 and #2) and from jri1401#15 hiPSCs using shATPIF1 #1. Unless otherwise indicated, 1383D6-derived IF1-KD cells were used throughout the study, and use of jri1401#15-derived IF1-KD cells is indicated in the figure legends. Stable IF1-OE cells were generated from 1383D6 hiPSCs. NFATc3 was transiently overexpressed in jri1401#15 hiPSCs, and the cells were harvested 3 days after transduction.

### Induction of Differentiation

hiPSCs were seeded at a density of 3 × 10⁵ cells per well in a 12-well plate in StemFit supplemented with 10 µM Y-27632 for 24 hours. Afterward, the cells were washed with DMEM/F12 (Thermo Fisher, #11330032) to remove Y-27632, and the medium was replaced with an induction medium. For mesendoderm induction, the cells were cultured in RPMI 1640 (Thermo Fisher, #11875093) supplemented with 1 × GlutaMAX (Thermo Fisher, #35050061) and 5 µM CHIR99021 (Cayman Chemical, #13122) (RPMI/CHIR medium; (Lam et al., 2014)) for up to 33 hours. For definitive endoderm induction, the cells were cultured in RPMI/CHIR medium for 24 hours, followed by culture in RPMI 1640 medium supplemented with 100 ng/mL Activin A (APRO Science, #GF-001) for an additional 24 hours. For neural progenitor cell induction, the cells were cultured in StemFit supplemented with 10 µM SB431542 (Sigma-Aldrich, #S4317-5MG) and 0.5 µM LDN193189 (Stemgent, #04-0074) (SF/2Si medium; (Townshend et al., 2020)) for 9 days with daily medium changes.

### Reverse transcription quantitative PCR (RT-qPCR)

For standard experiments, total RNA was extracted by a phenol-based method using TRIzol Reagent (Thermo Fisher, #15596026) or RNAzol RT (Molecular Research Center, #RN190). For time-course sample collection, total RNA was extracted using the column-based RNeasy Plus Mini Kit (QIAGEN, #74134). All extractions were performed according to the manufacturer’s instructions. Complementary DNA (cDNA) was synthesized from total RNA using the PrimeScript RT reagent kit (Takara, #RR037A). RT-qPCR was performed on LightCycler 2.0 (Roche) or StepOnePlus Real-Time PCR System (Applied Biosystems) using TB Green Premix Ex Taq II (Takara, #RR820A) and specific primer sets (Table S2). All reactions were run on the same instrument within each experiment, and data were not compared across different instruments. Whenever possible, primers were designed to span at least one intron, minimizing amplification of genomic DNA. Relative mRNA levels were calculated by the comparative Ct method using amplification efficiencies determined from standard curves, and normalized to β-actin (*ACTB*), glyceraldehyde-3-phosphate dehydrogenase (*GAPDH*), or ATP synthase peripheral stalk-membrane subunit b (*ATP5PB*). For each experiment, the reference gene was selected after confirming that its expression did not differ among the groups.

### Western blotting

For whole cell lysates, samples were collected using lysis buffer containing 1% TritonX-100 as previously described (Takamatsu et al., 2017). For nuclear/cytoplasmic fraction separations, separated samples were prepared using NE-PER Nuclear and Cytoplasmic Extraction Reagents (ThermoFisher, #78833) following the manufacturer’s protocol. Protein concentrations were measured using the Bradford assay (Bio-Rad, #5000006). Equal amounts of protein (10–25 µg) were separated on in-house SDS-PAGE gels of 8–15% acrylamide depending on the target molecular weight. For IF1 detection, either a 15% gel using Okajima’s low-molecular-weight protein separation method (Okajima et al., 1993) or a 15–20% gradient tris-tricine gel SuperSep Ace (FUJIFILM Wako, #198-15301) with Tricine Running Buffer (FUJIFILM Wako, #200-17071) was used. Proteins were transferred to nitrocellulose or PVDF membranes and blocked with Blocking One (Nacalai Tesque, #03953-95) for 30 minutes at room temperature. Membranes were incubated with primary antibodies overnight at 4℃, followed by incubation with HRP-conjugated secondary antibodies for 1 hour at room temperature. Primary and secondary antibody dilutions are listed in Table S3. Signals were visualized using Chemi-Lumi One Ultra (Nacalai Tesque, #11644-40), ECL Prime (Cytiva, #RPN2232), or ECL Start (Cytiva, #RPN3243) and quantified using CS analyzer software (ATTO, #2110030) or Fiji software (ImageJ) (Schindelin et al., 2012). Band intensities were normalized to β-actin (whole cell lysates), Lamin B1 (nuclear fraction), or α-tubulin (cytoplasmic fraction) as internal controls, unless otherwise specified.

### Immunocytochemistry

Cells were fixed for 10 min with 3.8% formaldehyde in PBS, diluted from Formaldehyde Solution (ca. 38% formaldehyde, 5–10% methanol; FUJIFILM Wako, #064-00401), and washed with PBS containing 10 mM glycine. Unless otherwise indicated, cells were permeabilized with 0.1% Triton X-100 in PBS for 10 min and blocked with 3% bovine serum albumin (BSA; Sigma-Aldrich, #A2153) in PBS for 30 min. Primary antibodies were diluted in 1% BSA in PBS and incubated at 4°C overnight. After washing with 0.1% BSA in PBS, cells were incubated with secondary antibodies diluted in 1% BSA in PBS for 45 min at 30°C. For NANOG staining, cells were permeabilized with 0.5% Triton X-100 in PBS for 10 min, blocked with G-Block (GenoStaff, #GB-01) at room temperature for 10 min, and incubated with primary antibodies diluted in 1% G-Block in PBS at 4°C for 3 days. Secondary antibodies were also diluted in 1% G-Block in PBS. Nuclei were counterstained with 0.5 µg/mL DAPI (Thermo Fisher, #D1306) during secondary antibody incubation. Primary and secondary antibody dilutions are listed in Table S3. Fluorescence images were acquired using an inverted fluorescence phase-contrast microscope (Keyence, BZ-X810). For each experiment, 10 fields were analyzed with five ROIs per field, and the mean N-cadherin/E-cadherin intensity ratio across all ROIs was used as the representative value.

### RNA sequencing and enrichment analysis

RNA was isolated using the RNeasy Plus Mini Kit (Qiagen). Libraries were prepared from total RNA of shScramble and shATPIF1 #1 hiPSCs, with three independent replicate samples per line, using the TruSeq Stranded mRNA Library Preparation Kit (Illumina). Sequencing was performed on the NovaSeq 6000 platform (Illumina), generating 101-bp paired-end reads. Total reads ranged from 41,268,506 to 55,895,142 per sample. Reads were preprocessed using fastp (Chen et al., 2018). Reads were aligned to the hg19 reference genome using STAR (v2.6.1d) (Dobin et al., 2013), following our laboratory’s established analysis pipeline via bcbio-nextgen (v1.2.9) (Chapman et al., 2021). In parallel, transcript abundance was quantified using Salmon (Patro et al., 2017) and summarized to gene-level TPM values using tximport (Soneson et al., 2015). Gene-level TPM values were used for visualization of expression heatmaps. Read counts obtained by featureCounts (Liao et al., 2014) were imported into iDEP (v2.4.4) (Ge et al., 2018), an R-based integrated differential expression and pathway analysis platform, for clustering, differential gene expression analysis, and enrichment analysis. Genes with CPM < 0.5 in all samples were excluded from further analysis. For PCA, count data were transformed by the edgeR log2(CPM + 4) method. Differential gene expression analysis was performed on the filtered, untransformed count data using DESeq2 (Love et al., 2014). P values were adjusted for multiple testing by the Benjamini–Hochberg procedure, and the resulting adjusted P values are reported as the false discovery rate (FDR) throughout. Genes with a false discovery rate (FDR) < 0.1 and an absolute fold change > 2 (|log2 fold change| > 1) were considered differentially expressed (Table S1). DEGs were subjected to GO biological process (Ashburner et al., 2000) and KEGG pathway (Kanehisa et al., 2021) enrichment analyses. Fold enrichment (observed/expected proportion of DEGs per pathway) was used as a measure of effect size. Terms passing an FDR cutoff of 0.05 were ranked by FDR, and the numbers of terms shown are given in the Figure 3 legend.

### Mitochondrial isolation

hiPSCs were seeded at a density of 1 × 10^5^ cells per well in 6-well plates and harvested on day 4. HeLa cells and TIG-1 fibroblasts were seeded at a density of 2 × 10^6^ and 1 × 10^6^ cells per 10-cm dish, and harvested on days 2 and 3, respectively. Cells were washed with PBS and detached using 0.5× TrypLE Select for hiPSCs, 0.05% trypsin-EDTA for HeLa cells, and 0.25% trypsin-EDTA for TIG-1 fibroblasts. The cells were centrifuged and the cell pellets were resuspended in 1 mL of ice-cold homogenization buffer containing 10 mM HEPES-KOH (pH 7.4), 220 mM mannitol, 70 mM sucrose, 10 μg/mL leupeptin and 10 μg/mL aprotinin. After incubation on ice for 15 min, the cells were homogenized by 30 passages through a 27-gauge needle. The homogenate was centrifuged at 900 × g for 5 min at 4°C to remove nuclei and cell debris. The supernatant was centrifuged at 5,000 × g for 10 min at 4°C to pellet the mitochondrial fraction. The pellet was resuspended in homogenization buffer and centrifuged again at 5,000 × g for 10 min at 4°C. The final mitochondrial pellet was resuspended in 25–40 μL of storage buffer containing 10 mM HEPES-KOH (pH 7.4) and 300 mM trehalose. Isolated mitochondria were stored at −80°C for up to 1 month until ATP hydrolysis activity measurement.

### ATP hydrolysis activity assay

Mitochondrial ATP hydrolysis activity was measured based on a previously described protocol with minor modifications (García-Bermúdez et al., 2016). Isolated mitochondria were subjected to three cycles of freeze-thaw using liquid nitrogen and a 37°C water bath to permeabilize the mitochondrial membrane. The mitochondrial protein concentrations were determined by BCA assay and adjusted to 1 μg/μl in the storage buffer described above.

The ATP hydrolysis activity assay was performed in a 96-well plate format. Each well contained 80 μl of reaction buffer with the following composition: 50 mM Tris-HCl (pH 8.0), 20 mM MgCl₂, 50 mM KCl, 5 mg/ml BSA, 5 μM FCCP, 1 μM antimycin A, 10 mM phosphoenolpyruvic acid monocyclohexylammonium salt (Tokyo Chemical Industry, #P0758), 2.5 mM adenosine 5’-triphosphate disodium salt hydrate (Tokyo Chemical Industry, #A0157), 1 mM β-nicotinamide adenine dinucleotide, reduced, dipotassium salt (NADH; Sigma-Aldrich, #N4505), 4 U/well L-lactate dehydrogenase (FUJIFILM Wako, #30052721), and 4 U/well pyruvate kinase (Sigma-Aldrich, #P7768). The reaction was initiated by the addition of 20 μL of mitochondrial suspension (20 μg mitochondrial protein).

Absorbance at 340 nm was measured using a microplate reader (Tecan Infinite 200 PRO M Plex) every 30 s for 10 min. Data from the initial 5 min were used to calculate the ATP hydrolysis rate. The decrease in NADH concentration was calculated using the Beer–Lambert equation, assuming an optical path length of 1 cm:

Δ[NADH] = ΔA₃₄₀ / (ε₃₄₀ × l)

where Δ[NADH] is the decrease in NADH concentration, ΔA₃₄₀ is the decrease in absorbance at 340 nm, ε₃₄₀ is the molar extinction coefficient of NADH at 340 nm (6.22 × 10³ M⁻¹ cm⁻¹), and l is the assumed optical path length. Apparent ATP hydrolysis activity was calculated from the rate of NADH oxidation, converted to nmol/min, and normalized to mitochondrial protein amount. Relative comparisons were made only between samples measured on the same plate under identical assay conditions. Since oligomycin was not routinely used for background subtraction, the values represent total mitochondrial fraction-associated ATP hydrolysis activity, which may include F1Fo-ATP synthase-associated ATP hydrolysis together with oligomycin-insensitive activities.

### Cell proliferation assay

For assessment of hiPSC proliferation, cells were seeded at 2.5 × 10⁴ cells/well in eight wells of a 24-well plate on Day 0, with two wells assigned to each time point (Day 1–Day 4). On the corresponding day, the cells were detached with 0.5× TrypLE Select, and viable cells were counted using a hemocytometer with trypan blue exclusion. The average cell count from the two wells was used as a single representative value. Data were obtained from three independent experiments.

### Quantification of ΔΨm, ROS, and mass

For quantification of mitochondrial parameters, 1 × 10⁴ hiPSCs per well were seeded on iMatrix-511-precoated (0.33 µg/well) black-walled, clear-bottom 96-well microplates (PerkinElmer, #6005182) three days prior to the assay. Thirty minutes before staining, the medium was replaced with fresh StemFit. For ΔΨm measurements, respiratory chain inhibitors were added to final concentrations of 10 µM OL (Sigma-Aldrich, #75351), 2 µM Rot (Adipogen Life Sciences, #AG-CN2-0516-G0001), 2 µM AA (Enzo Life Sciences, ALX-380-075-M005), and 50 µM FCCP (Abcam, #ab120081), and the following five conditions were tested: no inhibitor (basal), OL, Rot/AA, FCCP, and FCCP + OL. Cells were stained by replacing half the medium with fresh StemFit containing one of the following fluorescent probes at 2× final concentration and incubated at 37°C for 30 min: 2 µM JC-1 (Dojindo, #349-09401; JC-1 MitoMP Detection Kit), 500 nM MitoSOX Red (MSR; Thermo Fisher, #M36008), or 100 nM MitoTracker Green FM (MTG; Thermo Fisher, #M7514). After staining, the cells were washed twice with DMEM/F12 and placed in the appropriate buffer: Imaging Buffer supplied with the Dojindo kit for JC-1 or HEPES buffer (20 mM HEPES, 153 mM NaCl, 5 mM KCl, 5 mM glucose, pH 7.4) for MSR and MTG.

All fluorescence was measured at room temperature in top-read mode using a fluorescence microplate reader (Tecan, Infinite 200 Pro). A black adhesive seal was applied to the plate bottom to minimize background fluorescence. Excitation/emission settings were as follows: JC-1 (monomer: ex 485/em 530 nm, aggregate: ex 535/em 590 nm), MSR (ex 510/em 610 nm), and MTG (ex 485/em 530 nm). For nuclear normalization of the MTG signal, Hoechst 33342 (1 µg/mL) was added to MTG-treated wells and incubated for 20 min at 37°C, and fluorescence was measured (ex 350/em 461 nm) in HEPES buffer. ΔΨm, mtROS, and mitochondrial mass were calculated as the JC-1 aggregate/monomer ratio, MSR/MTG ratio, and MTG/Hoechst fluorescence intensity ratio, respectively.

### Imaging of ΔΨm

To assess mitochondrial membrane potential, 2 × 10⁵ hiPSCs were seeded on 35-mm dishes (day 0) and imaged on day 4. Thirty minutes prior to JC-1 staining, the culture medium was replaced with fresh medium. JC-1 was then added to a final concentration of 2 µM, and cells were incubated for 30 min at 37°C. After incubation, cells were washed twice with DMEM/F12 and transferred to 1× Imaging Buffer. Fluorescence images were acquired using an inverted fluorescence microscope (Olympus IX81) equipped with a xenon light source, a 20× objective lens, a DP74 color camera, and a temperature-controlled chamber. JC-1 monomers and aggregates were detected using the filter sets U-MNIBA3 (ex 470–495/ em 510–550 nm) and U-MWIY2 (ex 545–580/ em >610 nm), respectively. Fluorescence intensities were quantified using Fiji (ImageJ, NIH) (Schindelin *et al*., 2012), and ΔΨm was calculated as the ratio of JC-1 aggregate to monomer fluorescence.

### Measurement of intracellular ATP levels

Intracellular ATP levels were measured using ATP Assay Kit-Luminescence (Toyo B-Net, IC2-100) according to the manufacturer’s protocol. hiPSCs were seeded at a density of 5 × 10^4^ cells per well in 48-well plates and cultured for 2 days. On the day of measurement, cells were treated for 1 hour in fresh StemFit with or without 2 µM OL. Following treatment, the cells were lysed using the extraction reagent included in the kit, and luminescence (relative light units, RLU) was promptly measured using a microplate reader (Corona Electric, SH-1000Lab). ATP concentrations were quantified from a standard curve generated with known concentrations of ATP. Total protein concentration was determined from the same lysates using the Pierce BCA Protein Assay Kit (Thermo Fisher Scientific, #23225), and ATP levels were normalized to the protein content of each sample.

### Measurement of oxygen consumption rate

OCR was measured using the Extracellular OCR Plate Assay Kit (Dojindo, #E297), which is based on a phosphorescent oxygen probe, according to the manufacturer’s instructions. Briefly, hiPSCs were seeded on a PerkinElmer 96-well plate as described above. The cells were incubated with the Oxygen Probe at 37 °C for 30 min and overlaid with mineral oil. Fluorescence intensities (ex 500 / em 650 nm) were then recorded kinetically every 10 minutes for 200 minutes at 37 °C in bottom-read mode using a multifunctional plate reader (Molecular Devices, SpectraMax iD5). In each experiment, three replicate wells on the same plate were read simultaneously, and their mean fluorescence intensity, corrected using the blank values, was used to derive a single OCR value. OCR (pmol/min) was calculated using the OCR Calculation Sheet ver. 2 provided by the manufacturer. Prior to the experiments, we confirmed that the addition of FCCP increased the OCR. For normalization, OCR values were divided by Hoechst fluorescence intensity.

### Determination of glucose consumption and lactate production

Glucose consumption and lactate production were determined using the Glucose Assay Kit-WST (Dojindo, #G264) and the Lactate Assay Kit-WST (Dojindo, #L256), respectively, according to the manufacturer’s protocols. Briefly, hiPSCs were seeded at a density of 0.5 × 10⁵ cells per well in 24-well plates on day 0 and maintained with daily medium change. On day 3, the culture medium was replaced with fresh StemFit, and supernatants were collected 3 hours later for analysis. Absorbance at 450 nm was measured using a microplate reader (Corona Electric, SH-1000Lab). Glucose consumption was calculated by subtracting the glucose concentration in the supernatant from that in the fresh StemFit. Lactate production was calculated as the difference between the lactate concentration in the supernatant and that in fresh StemFit. Cell numbers were manually counted using trypan blue exclusion, and the results were normalized to cell number.

### Electron microscopy analysis

hiPSCs were detached with 0.5 mM EDTA in PBS at 37°C for 7 minutes and seeded at a density of 5.0 × 10⁴ cells onto Cell Desk LF1 (Sumitomo Bakelite, #MS-92132) in 24-well plates, with iMatrix-511 added directly to the cell suspension. The cells on the Cell Desk LF1 were immersed in the fixative containing 2.5% (volume (v) / v) glutaraldehyde and 2.0% (weight (w) / v) paraformaldehyde in phosphate buffer (PB; 0.1M, pH 7.4) overnight at 4°C. After rinsing with PB, the cells were postfixed with 1% (w/v) OsO4 in PB for 2 hours at 4°C. After rinsing with distilled water, they were stained by 2% (w/v) uranium acetate aqueous solution overnight at 4°C and embedded in epoxy resin according to standard procedures. Ultra-thin sections were then observed under an electron microscope (JEOL Ltd., JEM-1400).

### Ca²⁺ imaging

hiPSCs were detached with 0.5 mM EDTA in PBS at 37 °C for 7 min and seeded at a density of 1 × 10^5^ cells per 35-mm dishes (ibidi, #81156) in suspension containing iMatrix-511 and cultured for 3 days. Cells were incubated in StemFit containing 5 µM CalRed™ R525/650 AM (AAT Bioquest, #20590) and 0.04% Pluronic F-127 at 37 °C for 30 min. After dye loading, cells were washed twice with DMEM/F12 and transferred to 1× Ca²⁺-free HEPES buffer. Fluorescence images were acquired at room temperature using an Evident FV4000 confocal laser scanning microscope with a 20×/0.80 objective. CalRed was excited at 488 nm, and emission was collected at 500–540 nm (green) and 650–710 nm (red). Acquisition settings were as follows: scan speed, 2.0 µs/pixel; image size, 1024 × 1024 pixels; confocal aperture, 200 µm (1.47 Airy units); and three z-sections. For each channel, the three z-sections were combined by maximum-intensity projection in ImageJ, and the green/red fluorescence ratio was calculated.

For measurement of basal intracellular Ca²⁺ levels, fluorescence images were acquired under Ca²⁺-free conditions, and the green/red fluorescence intensity ratio was calculated as an index of intracellular Ca²⁺ levels. For each experiment, the ratio was averaged across 5 ROIs in each of 5 fields from one dish to give a single representative value.

For SOCE measurements, cells were analyzed under three conditions: control, 50 µM FCCP, or 10 µM OL. Thirty minutes before CalRed loading, cells were treated with fresh medium containing the indicated inhibitor, which was maintained throughout dye loading. After transfer to Ca²⁺-free HEPES buffer, thapsigargin (TPG; FUJIFILM Wako, #209-17281) was added to a final concentration of 2 µM and cells were incubated for 10 min. CaCl₂ was then added to a final concentration of 2 mM, and imaging was started immediately. Time-lapse images were acquired every 20 s for 16 frames over 5 min. SOCE was expressed as ΔR/R₀, where R₀ represents the green/red ratio of the first frame acquired immediately after CaCl₂ addition. For each experiment, 5 ROIs in a single field from one dish were averaged per frame to give the representative trace.

### Reversibility assays

For pharmacological rescue experiments, IF1-KD cells were treated with 1 µM cyclosporin A (CsA; FUJIFILM Wako, #031-24931) or 30 µM Ru265 (Sigma-Aldrich, #SML2991) for 24 h before RNA collection. CsA and Ru265 were dissolved in DMSO and dH_2_O, respectively, and the corresponding vehicle-treated cells were used as controls. For IF1 genetic restoration, IF1-KD cells were transduced with the lentiviral IF1-OE vector described above at an MOI of 10. Cells were collected for RNA extraction 48 h after transduction.

## QUANTIFICATION AND STATISTICAL ANALYSIS

### Number of replicates

Throughout this study, n denotes the number of independent experiments, defined as experiments performed on separate days using cells seeded independently for each experiment. Technical replicates within a single experiment were averaged before analysis, and the mean of each independent experiment was treated as one data point.

### Statistical Analysis

Statistical analyses were performed using GraphPad Prism version 11.0.2 (GraphPad Software, San Diego, California, USA). Data are presented as mean ± SD unless otherwise indicated. Statistical tests were chosen based on the experimental design and are described in each figure legend. Comparisons between two groups were performed using paired or unpaired two-tailed t-tests, as appropriate. For datasets better described by a lognormal distribution, lognormal t-tests were used as indicated. For multiple group comparisons, one-way or two-way ANOVA followed by appropriate post hoc tests, including Dunnett’s or Bonferroni’s multiple comparisons tests, were used. When multiple t-tests were performed, *P* values were adjusted using the Holm method where indicated. A *P* value < 0.05 was considered statistically significant.

## ADDITIONAL RESOURCES

Not applicable

### Supplemental information

Document S1. Figures S1–S5 and Table S2 and S3

### Declaration of generative AI and AI-assisted technologies in the writing process

During the preparation of this work the authors used AI-assisted tools (ChatGPT, OpenAI; Claude Sonnet and Opus, Anthropic) in order to assist with translation of the manuscript from Japanese into English and to suggest reorganization of the Discussion section for clarity and flow. All AI-assisted text was reviewed, revised, and approved by the authors to ensure accuracy and clarity of meaning. The authors take full responsibility for the accuracy and integrity of the work.

## Key Resources Table

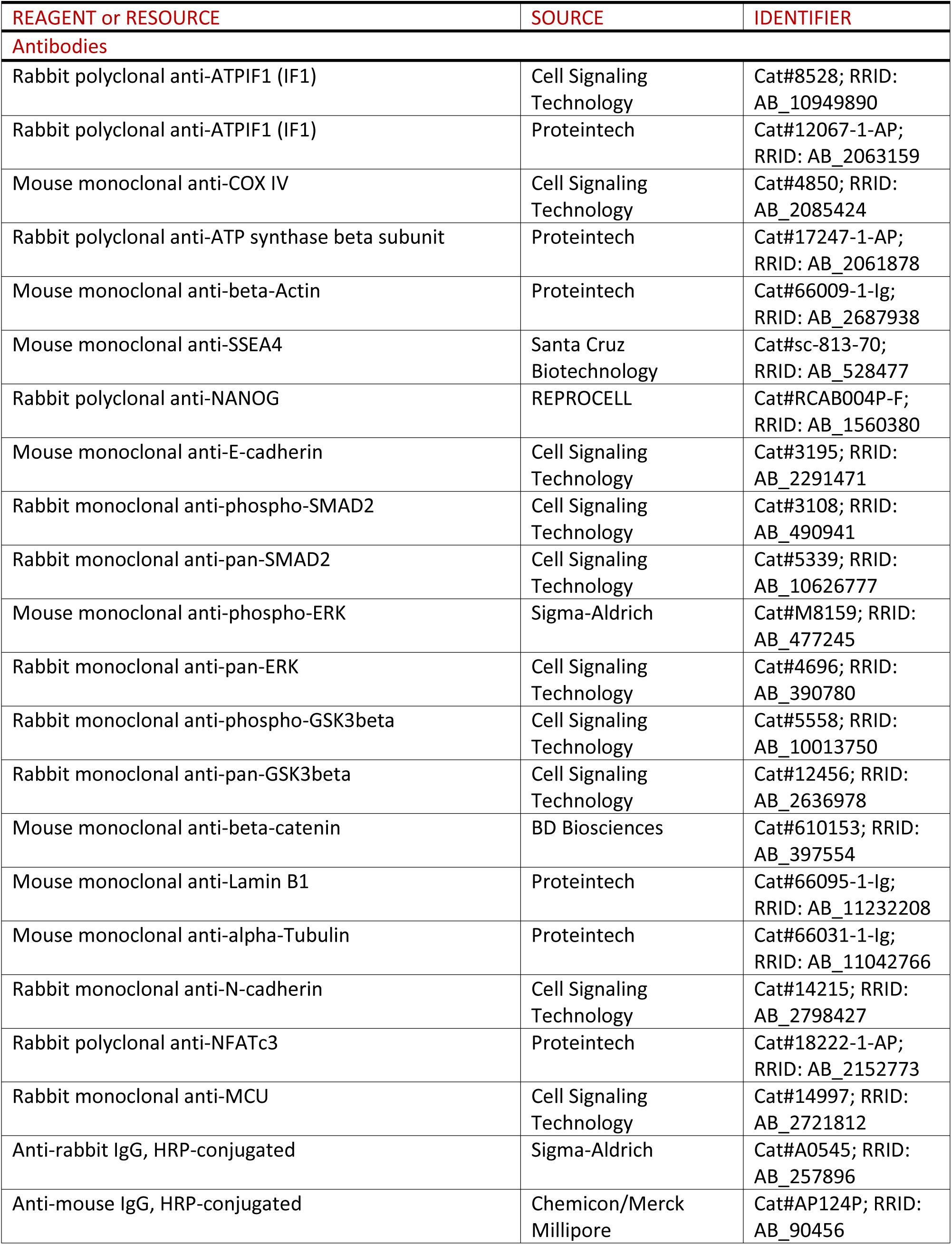
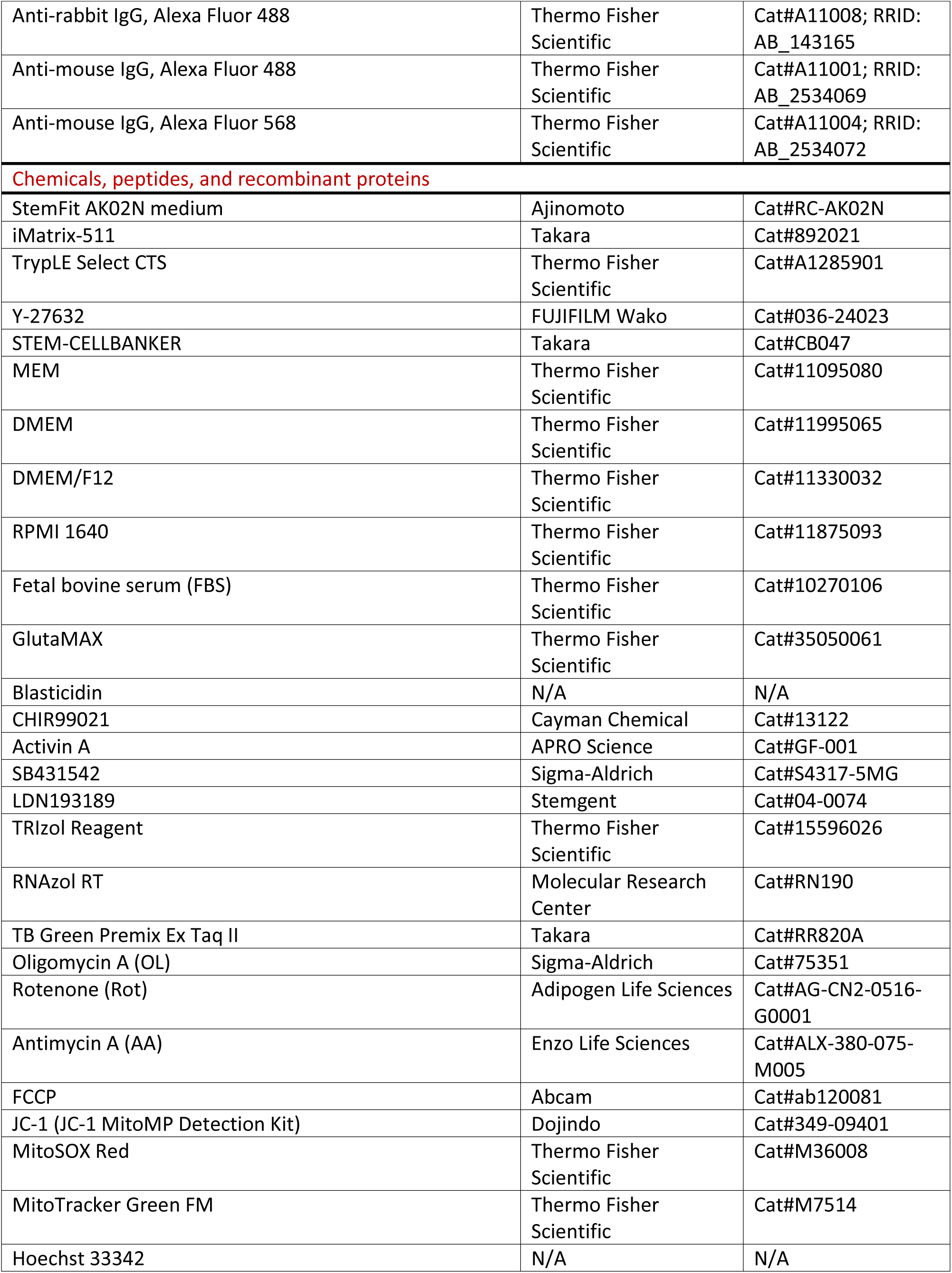
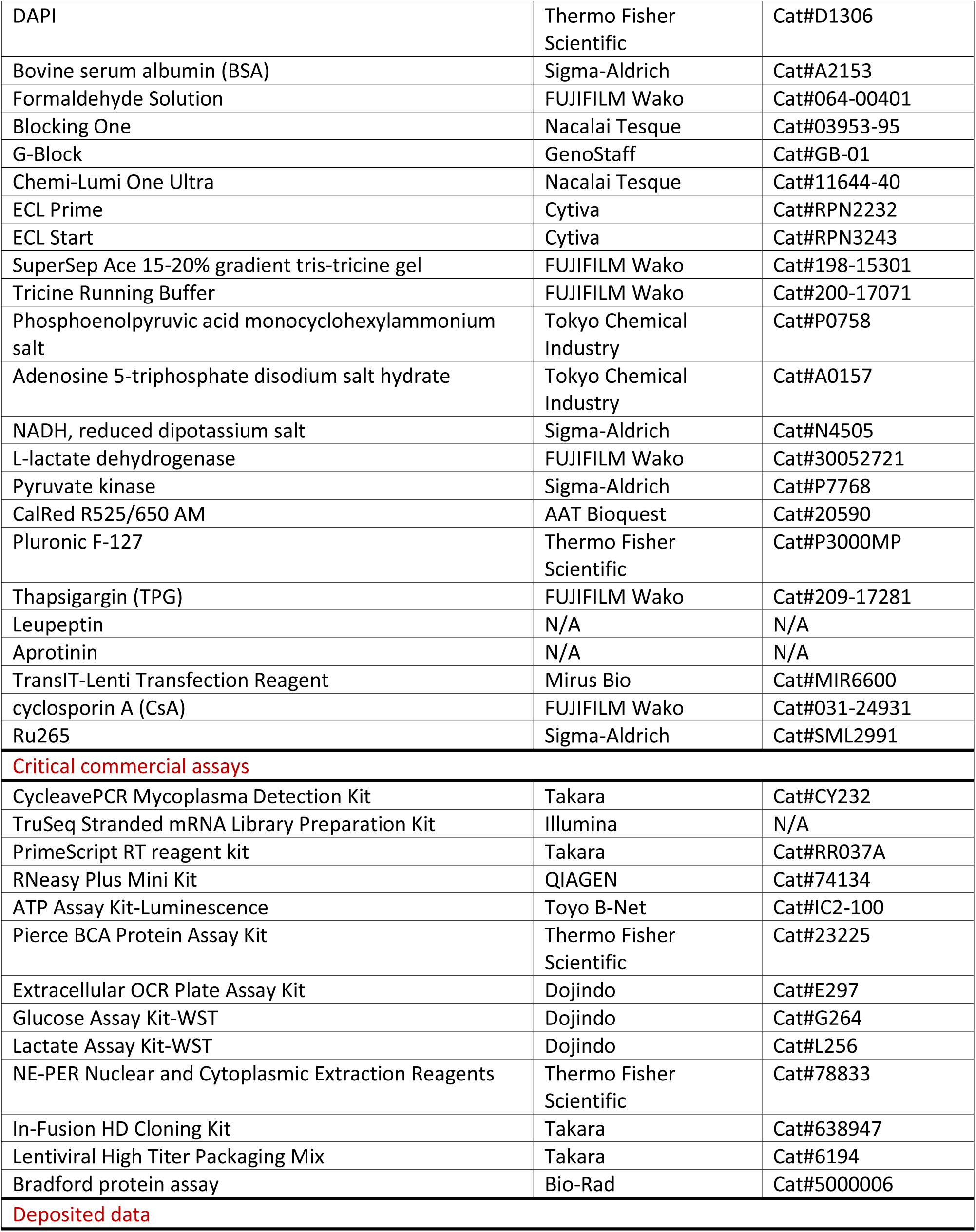
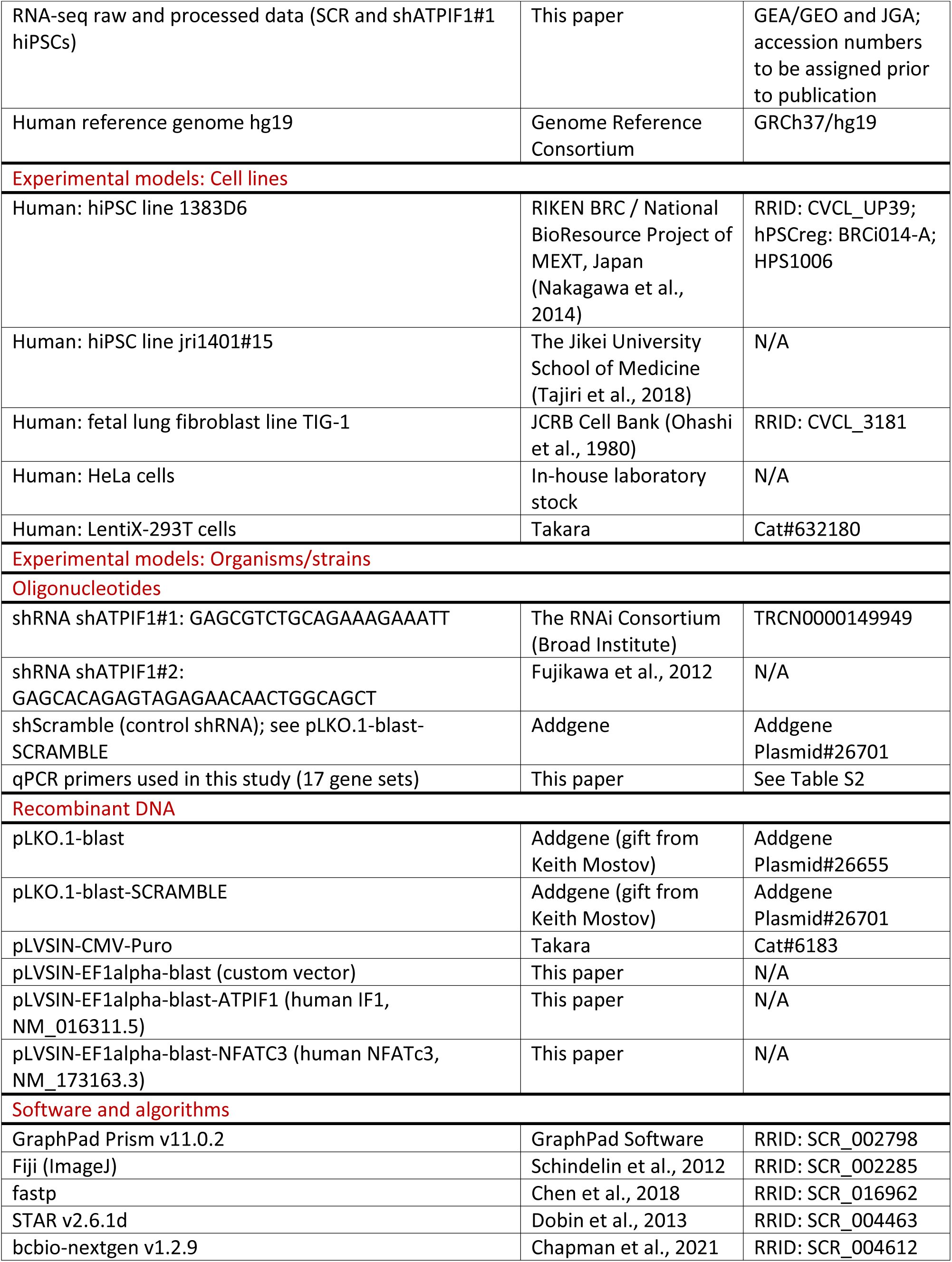
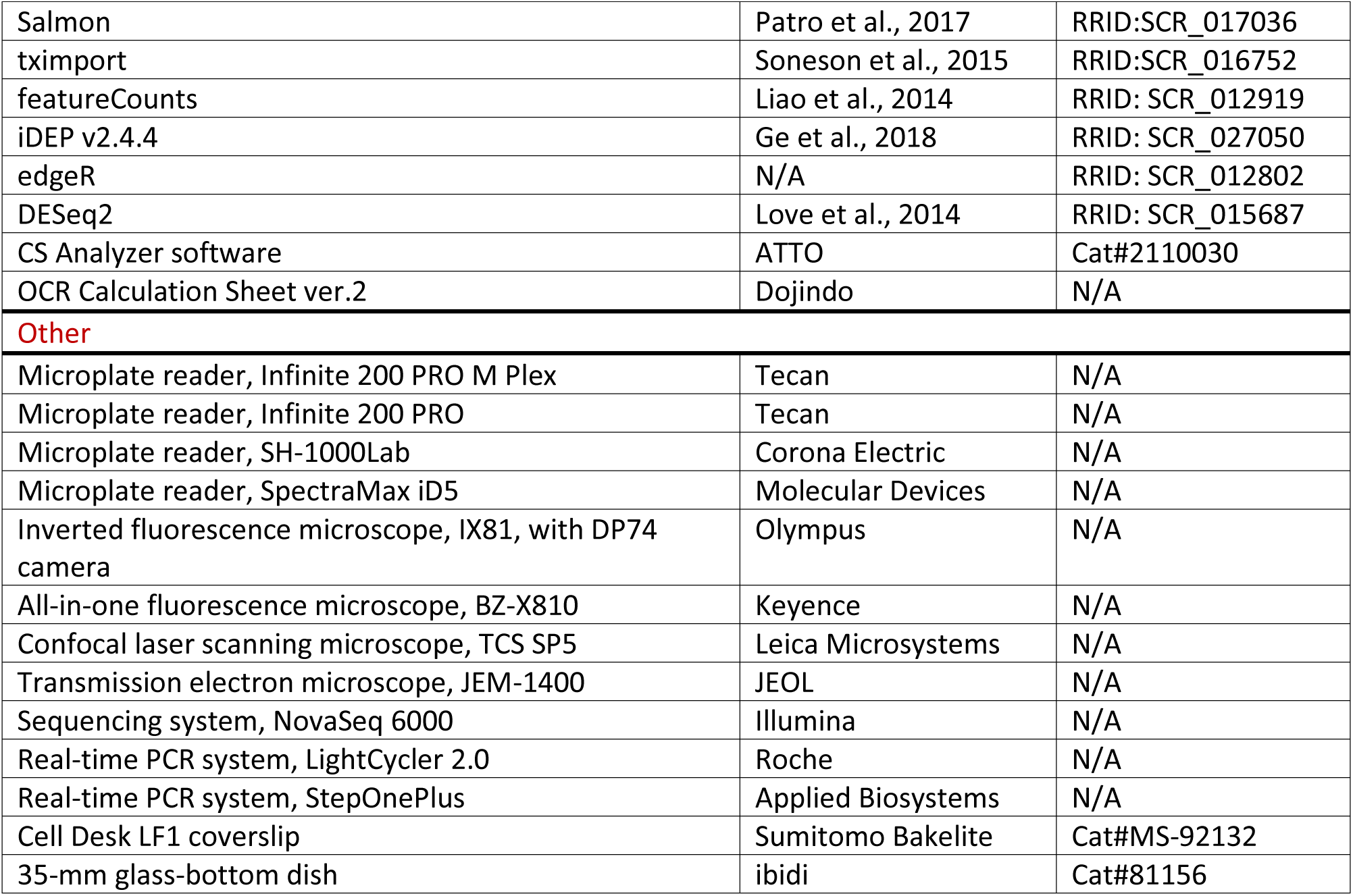

## Supplemental Figures

**Figure S1. Related to Figure 1.**
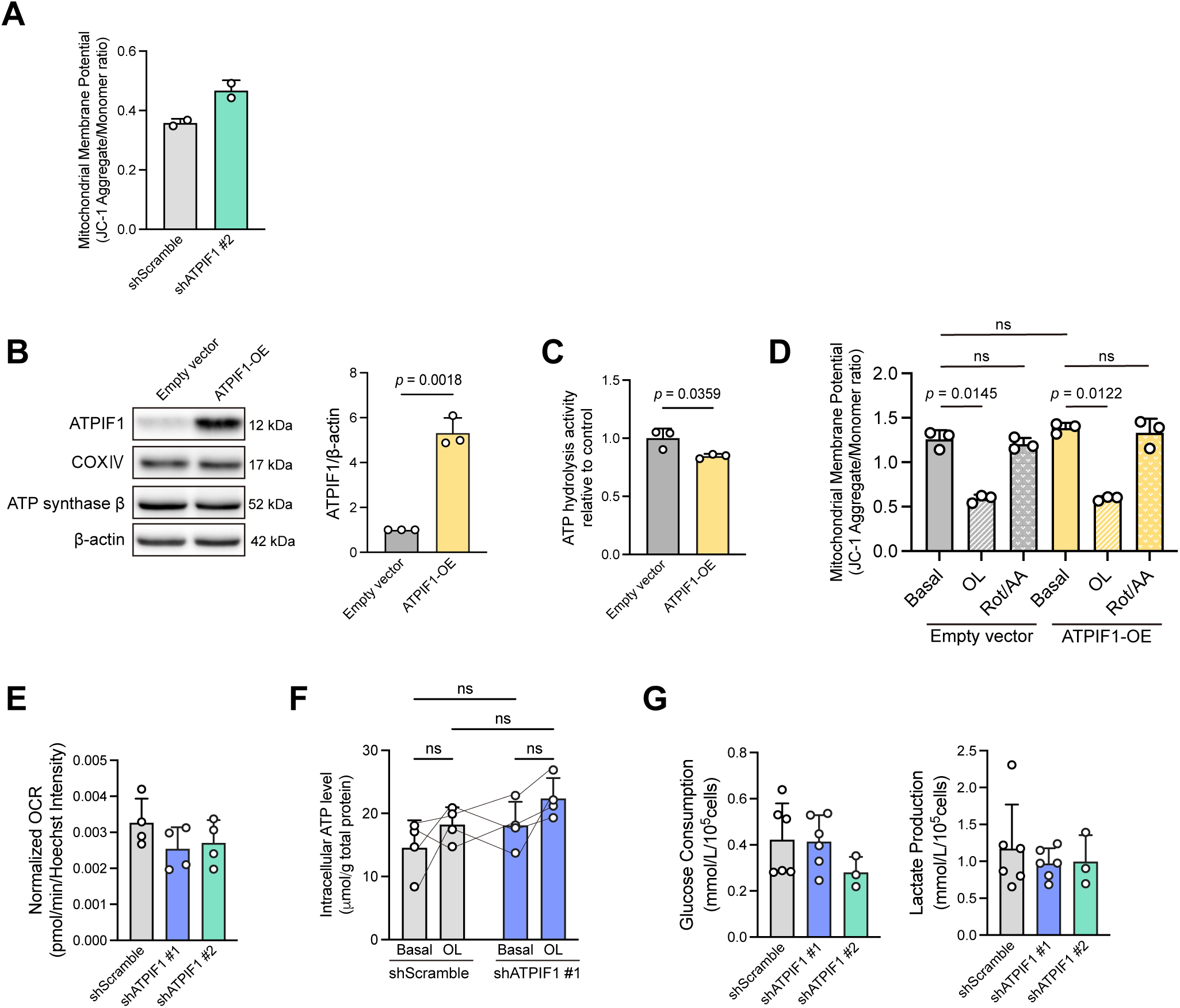
(A) ΔΨm, assessed by JC-1 fluorescence imaging, in shScramble and shATPIF1 #2 cells. Two independent experiments. (B–D) Empty vector and IF1-overexpressing (IF1-OE) cells. (B) Representative western blot images of IF1, COX IV, ATP synthase β, and β-actin, and quantification of relative IF1 protein levels. (C) ATP hydrolysis activity in isolated mitochondria, normalized to the mean of the control group (set to 1). (D) ΔΨm in cells treated with 10 μM OL or 2 μM each of Rot and AA (Rot/AA). Three independent experiments each. (E) Basal oxygen consumption rate (OCR) in shScramble, shATPIF1 #1, and shATPIF1 #2 cells. Data for shScramble and shATPIF1 #1 are the same as those shown in Figure 1I and are replotted for comparison. Four independent experiments. (F) Intracellular ATP levels, measured using a luciferase-based assay, in shScramble and shATPIF1 #1 cells in the presence or absence of 2 μM OL. Data obtained on the same experimental day are connected by lines. Four independent experiments. (G) Glucose consumption and lactate production in shScramble, shATPIF1 #1, and shATPIF1 #2 cells. Six independent experiments for shScramble and shATPIF1 #1, and three for shATPIF1 #2. Individual values with mean ± SD. Exact *P* values are shown when *P* < 0.05. Statistics: paired two-tailed t-test on log-transformed values (B–C), repeated-measures two-way ANOVA followed by Bonferroni correction (D, F), and repeated-measures one-way ANOVA followed by Dunnett’s multiple comparison test (E, G).

**Figure S2. Related to Figure 2.**
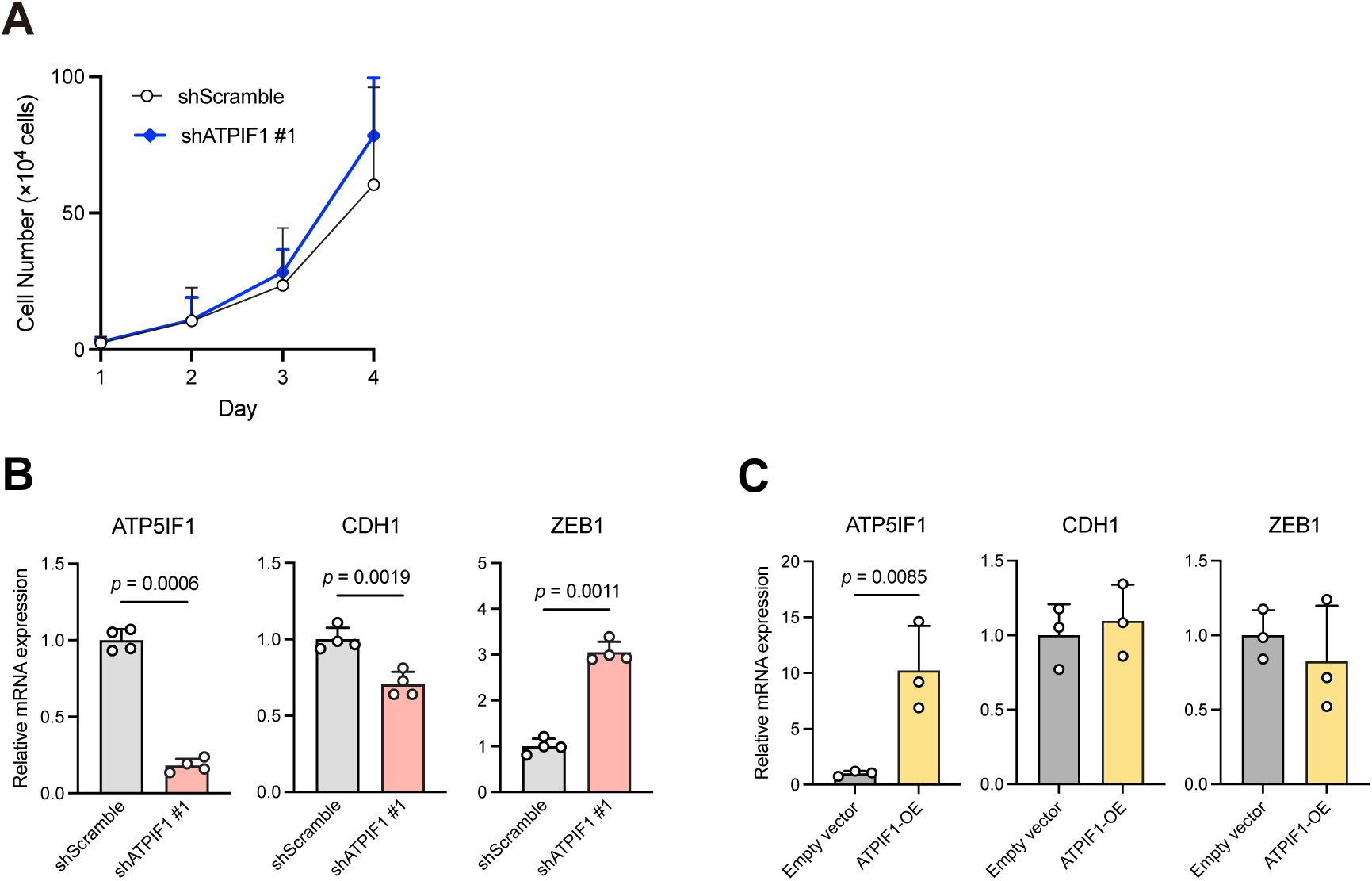
(A) Cell proliferation assay in shScramble and shATPIF1 #1 cells. Three independent experiments. (B) RT-qPCR of *ATP5IF1*, *CDH1* and *ZEB1* in shATPIF1 #1 cells generated from a different parental hiPSC line, jri1401#15, confirming the expression changes observed in shATPIF1 #1 cells derived from 1383D6. Four independent experiments. (C) RT-qPCR of *ATP5IF1*, *CDH1* and *ZEB1* in empty vector and IF1-OE cells. Three independent experiments. RT-qPCR data in (B) and (C) were normalized to *ACTB* and expressed relative to the mean of the corresponding control group (set to 1). Individual values with mean ± SD. Exact *P* values are shown when *P* < 0.05. Statistics: paired two-tailed t-test between shScramble and shATPIF1 #1 cells on the same day (A) and paired two-tailed t-test on log-transformed values (B, C).

**Figure S3. Related to Figure 3.**
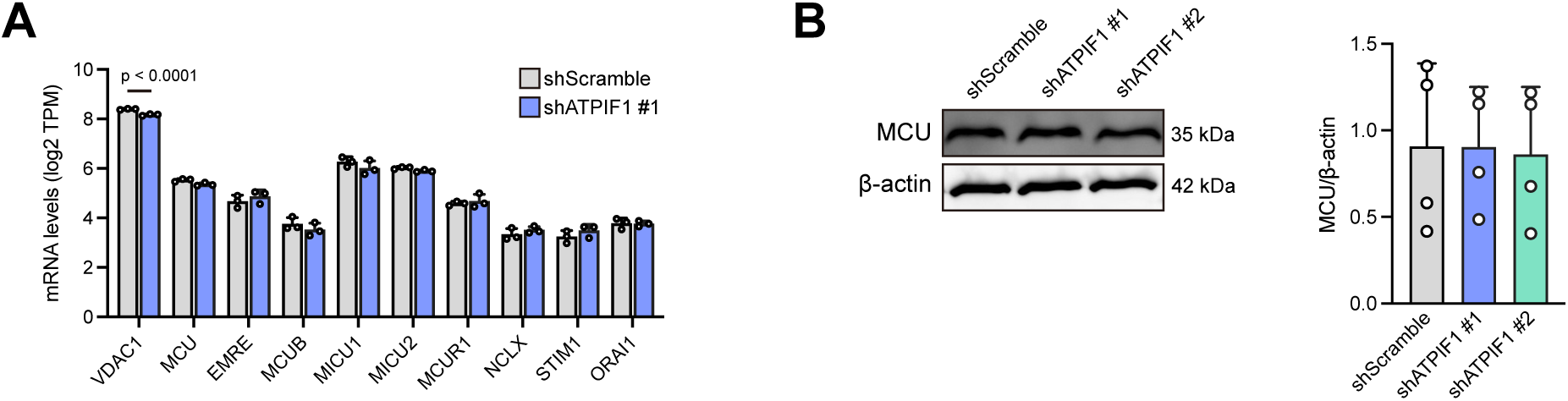
(A) Western blots and quantification of canonical EMT-related signaling in shScramble, shATPIF1 #1, and shATPIF1 #2 cells: phosphorylation of SMAD2, extracellular signal-regulated kinase 1/2 (ERK1/2), and glycogen synthase kinase-3β (GSK-3β), and nuclear translocation of β-catenin. Phosphorylation levels were normalized to the corresponding total protein, and nuclear β-catenin was normalized to Lamin B1. Three independent experiments. Individual values with mean ± SD. Exact *P* values are shown when *P* < 0.05. Statistics: repeated-measures one-way ANOVA followed by Dunnett’s multiple comparison test.

**Figure S4. Related to Figure 4.**
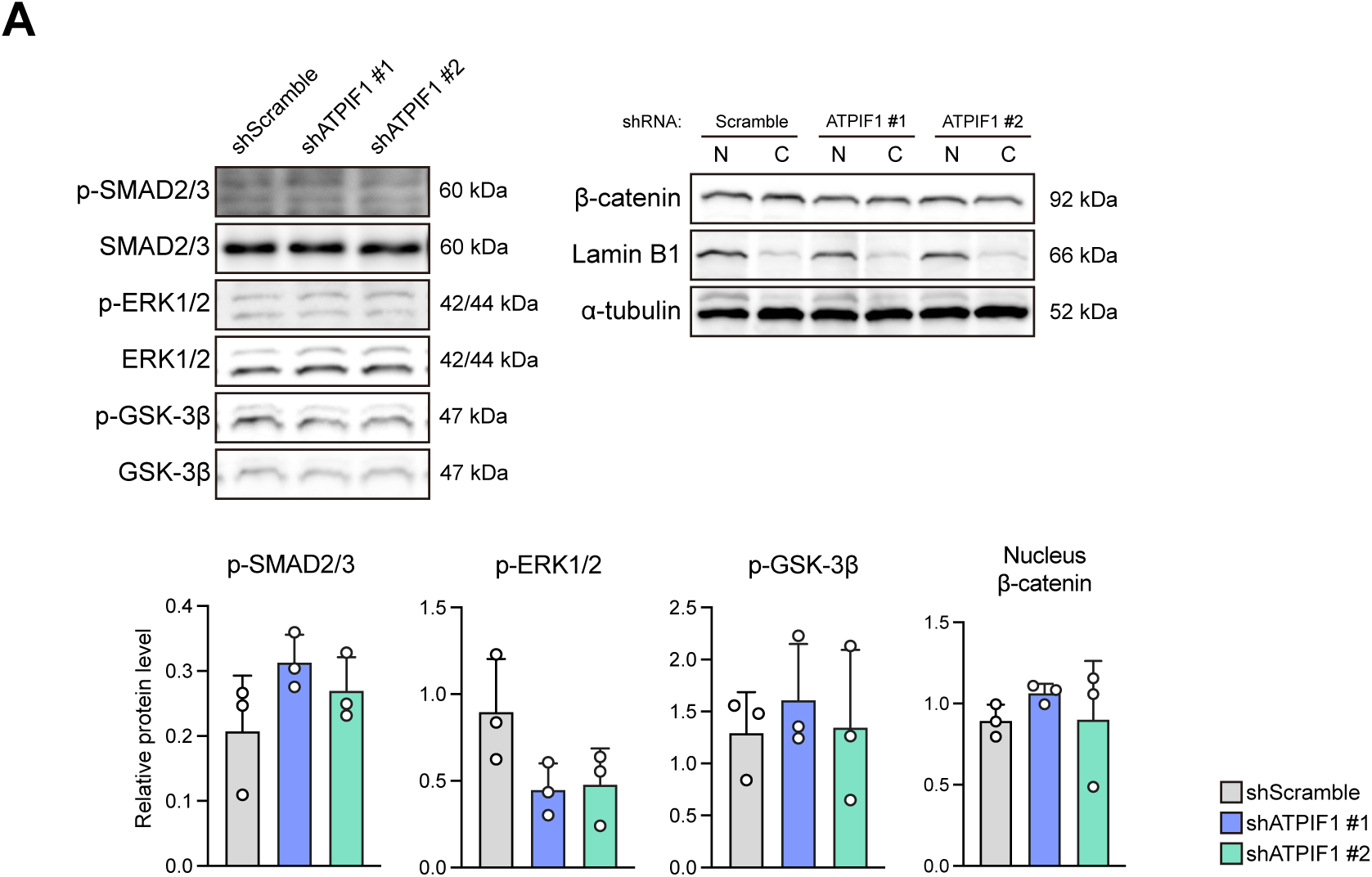
(A) mRNA levels (log2 TPM) of ten genes related to mitochondrial Ca²⁺ handling and SOCE, including *ORAI1*, *STIM1*, *MCU*, and voltage-dependent anion channel 1 (*VDAC1*), in shScramble and shATPIF1 #1 cells, from RNA sequencing. (B) Representative western blot images of MCU and quantification of relative MCU protein levels, normalized to β-actin, in shScramble, shATPIF1 #1, and shATPIF1 #2 cells. Four independent experiments. Individual values with mean ± SD. Exact *P* values are shown when *P* < 0.05. Statistics: DESeq2 adjusted *P* values (A) and repeated-measures one-way ANOVA followed by Dunnett’s multiple comparison test (B).

**Figure S5. Related to Figure 6.**
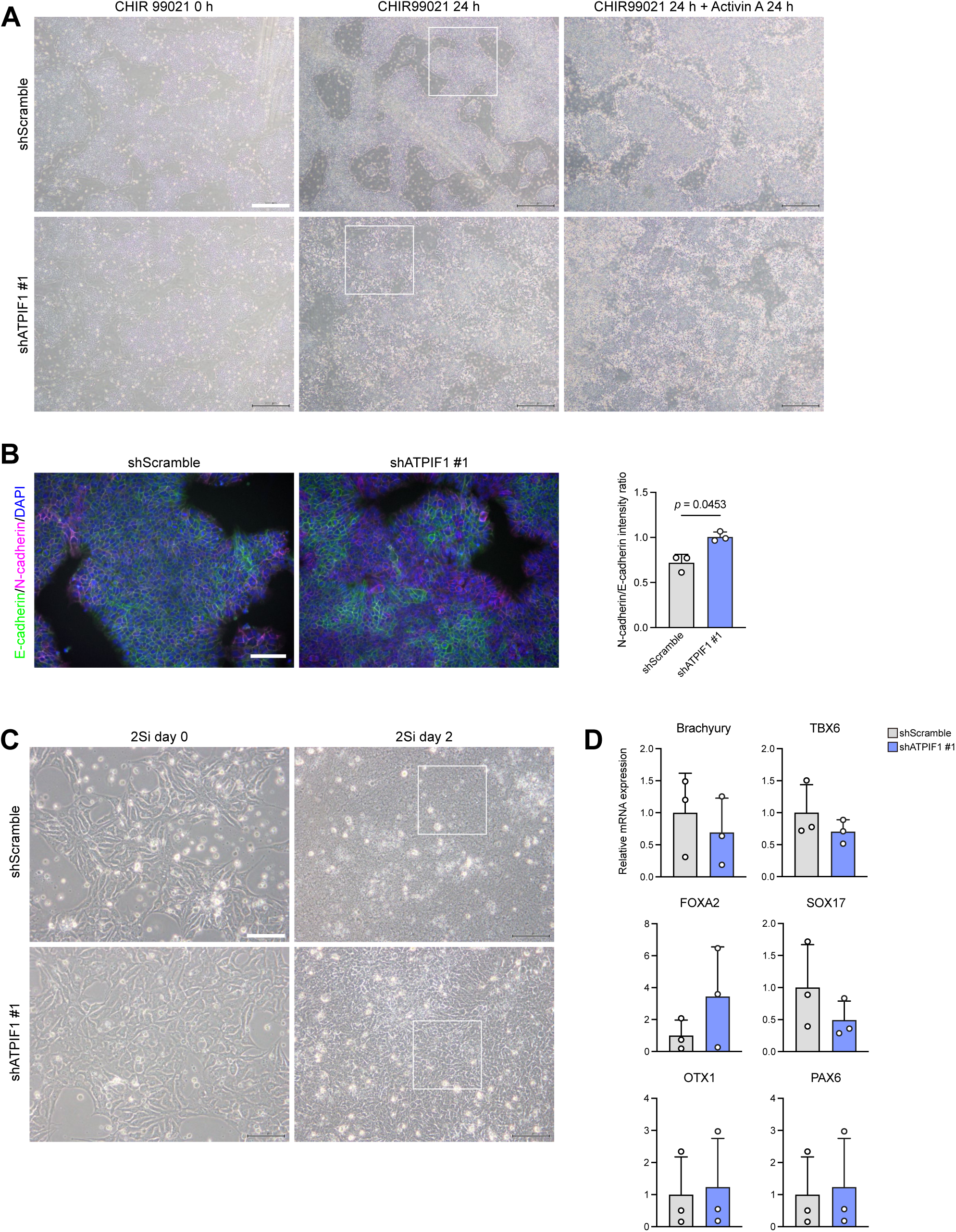
All panels compare shScramble and shATPIF1 #1 cells. Three independent experiments. (A) Phase-contrast images before and after mesendoderm and endoderm induction. Figure 6B is an excerpt of the boxed region (CHIR99021 24 h). Scale bar = 500 μm. (B) Representative immunocytochemistry images of E-cadherin (green), N-cadherin (magenta), and DAPI (blue) before mesendoderm induction (0 h), and quantification of the N-cadherin/E-cadherin intensity ratio. Scale bar = 100 μm. (C) Phase-contrast images before and after dual-SMAD inhibition (2Si) for neuroectoderm induction. Figure 6G is an excerpt of the boxed region (2Si day 2). Scale bar = 100 μm. (D) RT-qPCR of the mesendoderm/mesoderm markers *TBXT* (Brachyury) and *TBX6*, the definitive endodermal markers *SOX17* and *FOXA2*, and the neuroectodermal markers *OTX1* and *PAX6* in undifferentiated cells, normalized to *ACTB* and expressed relative to the mean of the control group (set to 1). Individual values with mean ± SD. Exact *P* values are shown when *P* < 0.05. Statistics: paired two-tailed t-test on log-transformed values (B, D).

**Table S1.** Differential expression analysis of shATPIF1 #1 versus shScramble cells. All genes retained after filtering are listed. baseMean, log2FC, and FDR were calculated with DESeq2; FDR denotes the Benjamini–Hochberg adjusted P value. Expression values for individual replicates (SCR_1–3, KD_1– 3) are shown as log2-transformed normalized counts. Genes with NA in the FDR column were excluded from statistical testing by the independent filtering procedure of DESeq2.

**Table S2.**
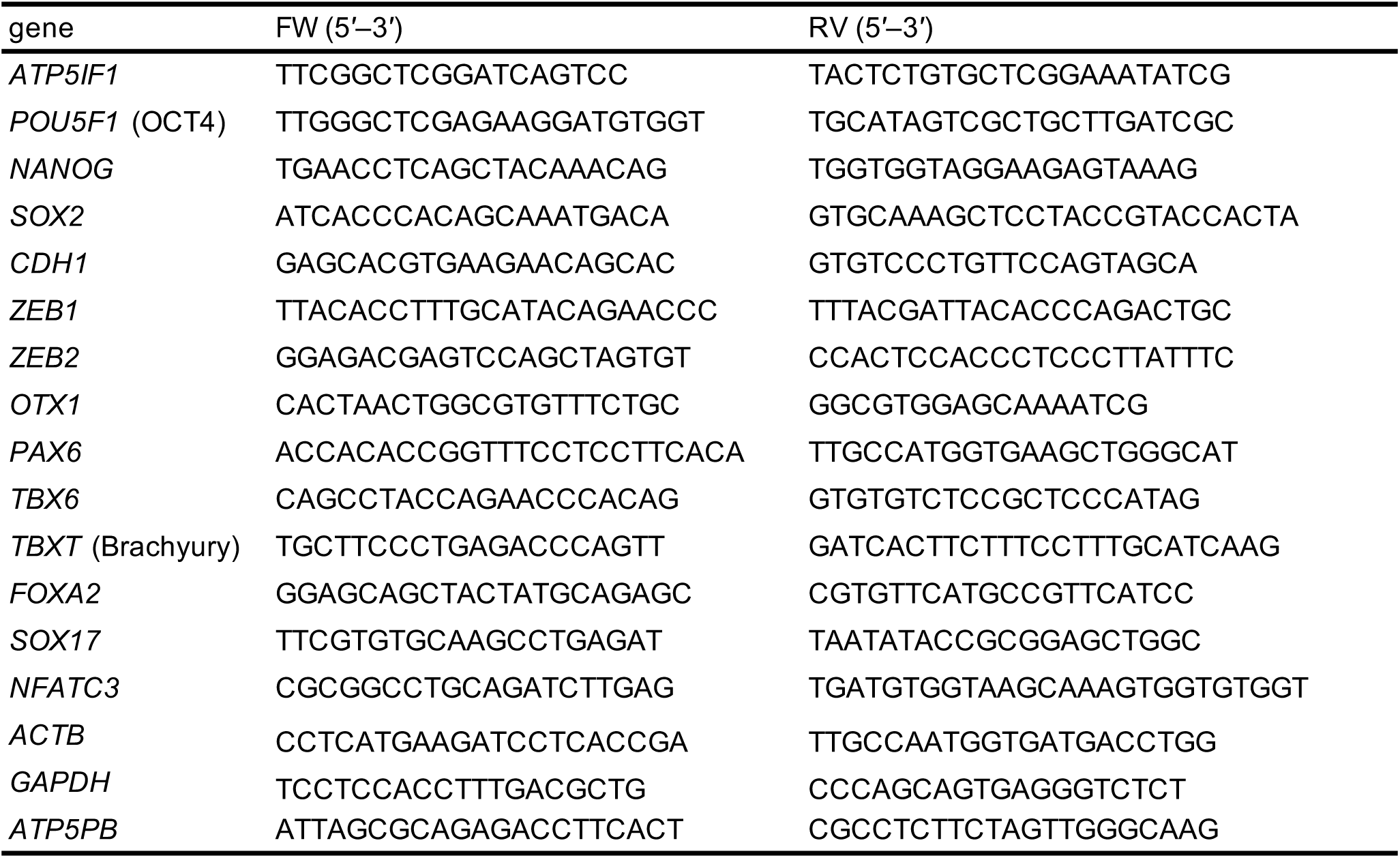
qPCR primer sequences used in this study.

**Table S3.**
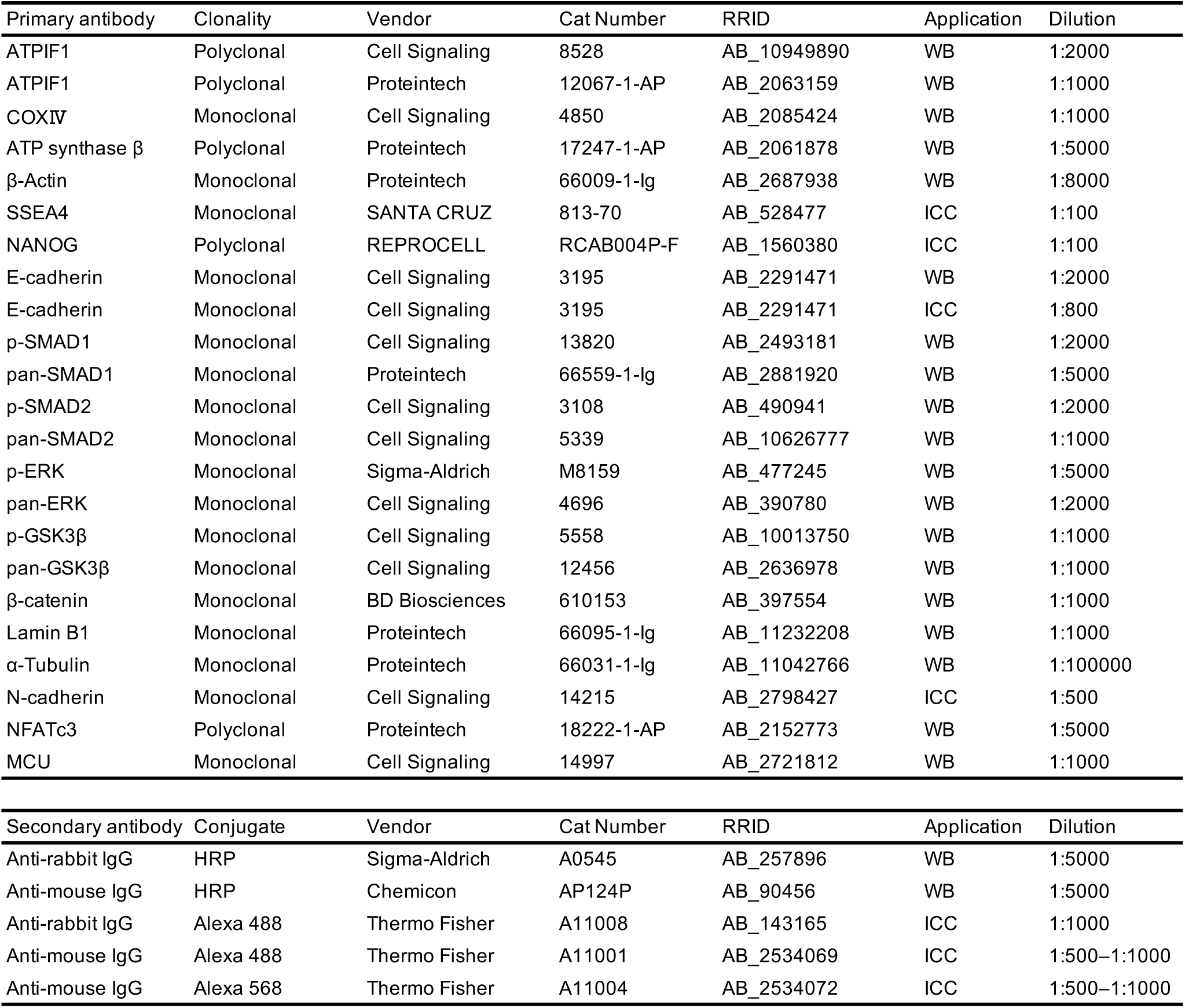
Antibodies dilution used in this study.

| Primary antibody | Clonality | Vendor | Cat Number | RRID | Application | Dilution |
| --- | --- | --- | --- | --- | --- | --- |
| ATPIF1 | Polyclonal | Cell Signaling | 8528 | AB_10949890 | WB | 1:2000 |
| ATPIF1 | Polyclonal | Proteintech | 12067-1-AP | AB_2063159 | WB | 1:1000 |
| COXIV | Monoclonal | Cell Signaling | 4850 | AB_2085424 | WB | 1:1000 |
| ATP synthase $\beta$ | Polyclonal | Proteintech | 17247-1-AP | AB_2061878 | WB | 1:5000 |
| β-Actin | Monoclonal | Proteintech | 66009-1-Ig | AB_2687938 | WB | 1:8000 |
| SSEA4 | Monoclonal | SANTA CRUZ | 813-70 | AB_528477 | ICC | 1:100 |
| NANOG | Polyclonal | REPROCELL | RCAB004P-F | AB_1560380 | ICC | 1:100 |
| E-cadherin | Monoclonal | Cell Signaling | 3195 | AB_2291471 | WB | 1:2000 |
| E-cadherin | Monoclonal | Cell Signaling | 3195 | AB_2291471 | ICC | 1:800 |
| p-SMAD1 | Monoclonal | Cell Signaling | 13820 | AB_2493181 | WB | 1:2000 |
| pan-SMAD1 | Monoclonal | Proteintech | 66559-1-Ig | AB_2881920 | WB | 1:5000 |
| p-SMAD2 | Monoclonal | Cell Signaling | 3108 | AB_490941 | WB | 1:2000 |
| pan-SMAD2 | Monoclonal | Cell Signaling | 5339 | AB_10626777 | WB | 1:1000 |
| p-ERK | Monoclonal | Sigma-Aldrich | M8159 | AB_477245 | WB | 1:5000 |
| pan-ERK | Monoclonal | Cell Signaling | 4696 | AB_390780 | WB | 1:2000 |
| p-GSK3β | Monoclonal | Cell Signaling | 5558 | AB_10013750 | WB | 1:1000 |
| pan-GSK3β | Monoclonal | Cell Signaling | 12456 | AB_2636978 | WB | 1:1000 |
| β-catenin | Monoclonal | BD Biosciences | 610153 | AB_397554 | WB | 1:1000 |
| Lamin B1 | Monoclonal | Proteintech | 66095-1-Ig | AB_11232208 | WB | 1:1000 |
| α-Tubulin | Monoclonal | Proteintech | 66031-1-Ig | AB_11042766 | WB | 1:100000 |
| N-cadherin | Monoclonal | Cell Signaling | 14215 | AB_2798427 | ICC | 1:500 |
| NFATc3 | Polyclonal | Proteintech | 18222-1-AP | AB_2152773 | WB | 1:5000 |
| MCU | Monoclonal | Cell Signaling | 14997 | AB_2721812 | WB | 1:1000 |

| Secondary antibody | Conjugate | Vendor | Cat Number | RRID | Application | Dilution |
| --- | --- | --- | --- | --- | --- | --- |
| Anti-rabbit IgG | HRP | Sigma-Aldrich | A0545 | AB_257896 | WB | 1:5000 |
| Anti-mouse IgG | HRP | Chemicon | AP124P | AB_90456 | WB | 1:5000 |
| Anti-rabbit IgG | Alexa 488 | Thermo Fisher | A11008 | AB_143165 | ICC | 1:1000 |
| Anti-mouse IgG | Alexa 488 | Thermo Fisher | A11001 | AB_2534069 | ICC | 1:500–1:1000 |
| Anti-mouse IgG | Alexa 568 | Thermo Fisher | A11004 | AB_2534072 | ICC | 1:500–1:1000 |

## Notes

### Competing Interest Statement

The authors have declared no competing interest.

### Summary of Updates

This version adds a new Results section describing the effects of IF1 re-expression and of pharmacological inhibition with cyclosporin A (calcineurin) and Ru265 (MCU) on the established EMT-like state, together with the corresponding figure. A supplementary file containing the RNA-seq data has also been added. Some preliminary data have been removed to focus the manuscript, the figures reorganized, and the text revised accordingly.

